# Microbiome Determinants and Physiological Effects of the Benzoate-Hippurate Microbial-Host Co-Metabolic Pathway

**DOI:** 10.1101/2019.12.15.876672

**Authors:** François Brial, Julien Chilloux, Trine Nielsen, Sara Vieira-Silva, Gwen Falony, Lesley Hoyles, Ana L. Neves, Andrea Rodriguez-Martinez, Ghiwa Ishac Mouawad, Nicolas Pons, Sofia Forslund, Emmanuelle Le Chatelier, Aurélie M. Le Lay, Jeremy K Nicholson, Torben Hansen, MetaHIT consortium, Karine Clément, Peer Bork, S. Dusko Ehrlich, Jeroen Raes, Oluf Pedersen, Dominique Gauguier, Marc-Emmanuel Dumas

## Abstract

**Objective:** Gut microbial products are involved in type 2 diabetes, obesity and insulin resistance. In particular, hippurate, a hepatic phase 2 conjugation product of microbial benzoate metabolism, has been associated with a healthy phenotype. This study aims to identify metagenomic determinants and test protective effects of hippurate.

**Design:** We profiled the urine metabolome by ^1^H Nuclear Magnetic Resonance (NMR) spectroscopy to derive associations with metagenomic sequences in 271 middle-aged Danish individuals to identify dietary patterns in which urine hippurate levels were associated with health benefits. We follow up with benzoate and hippurate infusion in mice to demonstrate causality on clinical phenotypes.

**Results:** In-depth analysis identifies that the urine hippurate concentration is associated with microbial gene richness, microbial functional redundancy as well as functional modules for microbial benzoate biosynthetic pathways across several enterotypes. Through dietary stratification, we identify a subset of study participants consuming a diet rich in saturated fat in which urine hippurate, independently of gene richness, accounts for links with metabolic health that we previously associated with gene richness. We then demonstrate causality *in vivo* through chronic subcutaneous infusions of hippurate or benzoate (20 nmol/day) resulting in improved glycemic control in mice fed a high-fat diet. Hippurate improved insulin secretion through increased β-cell mass and reduced liver inflammation and fibrosis, whereas benzoate treatment resulted in liver inflammation.

**Conclusion:** Our translational study shows that the benzoate-hippurate pathway brings a range of metabolic improvements in the context of high-fat diets, highlighting the potential of hippurate as a mediator of metabolic health.

## INTRODUCTION

The human obesity epidemic raises the risk of type 2 diabetes and cardiovascular disease. Dysbiosis of the gut microbiome is now recognised as a key feature of these disorders.[1] The microbiome collectively encodes up to 10 million different microbial genes.[2, 3] In particular, gene richness has been proposed as a marker of ecological diversity mirroring improvements in metabolic health.[4, 5] Although the microbiome directly impacts various biological processes of the host through production or degradation of a multitude of compounds, the vast majority of molecules involved in this chemical crosstalk remains elusive.[6–9]

There is growing evidence that hippurate, one of the most abundant microbial-host co-metabolites in human urine, is positively associated with metabolic health through inverse associations with blood pressure, fatty liver disease, visceral fat mass and Crohn’s disease.[10–14] Its microbial precursor, benzoate is taken up by organic anion transporter MCT2[15] and conjugated with glycine in the liver and kidney to form hippurate. We showed in a genetic cross between diabetic and normoglycemic rat strains that serum benzoate under host genome control.[16] Hippurate was recently shown to be associated both with microbiota diversity based on 16S rRNA gene sequencing[17] and with reduced risk of metabolic syndrome.[13]

However, there is a critical need for an in-depth characterization of the complex nutrition-microbiome-host interaction in the benzoate-hippurate pathway, in relation to: i) associations with enterotype, gene richness and functional redundancy, ii) population stratification according to nutritional patterns to identify patient sub-groups in which hippurate improves metabolic health, and iii) translational elucidation of these effects on host phenotypes *in vivo*. To address these specific points, we characterized the urinary metabolome and the fecal microbiome of 271 middle-aged non-diabetic Danish individuals from the MetaHIT study.[4] Through dietary stratification we delineate the complex interaction between dietary intake, microbiome and metabolome its impact on body weight, immune and metabolic markers, which we further confirm and characterise in a mouse model of diet-induced obesity and diabetes.

## METHODS

### Human subjects

All analyses were done on non-diabetic Danish individuals from the MetaHIT study (n=271),[4, 18] including the subset of 193 individuals who completed a validated food frequency questionnaire (FFQ).[19] The study was approved by the Ethical Committees of the Capital Region of Denmark (HC-2008-017 and H-15000306) and was in accordance with the principles of the Declaration of Helsinki. All individuals gave written informed consent before participating in the study. Sampling and clinical phenotyping were performed as described previously.[4, 18] In short, all study participants were recruited from the population-based Inter99 study.[20]. The study program consisted of two visits with approximately 14 days apart. At the first visit all participants were examined in the morning after an overnight fast. Blood sampling was performed at the fasting state, and urine was collected upon arrival at the centre. At the second visit, a Dual Energy X-Ray Absorptiometry (DXA) scan was performed. Serum glycine levels were previously assessed.[19] Estimated glomerular filtration rate (eGFR) was calculated with the CKD-EPI formula with age, gender and creatinine (μmol/L) and without ethnicity factor.[21]

### Dietary assessment

A subset of the study participants (n=193) completed a validated FFQ in order to obtain information on their habitual dietary habits.[22] The FFQ gathered dietary information from all meals during a day and the intake frequencies within the past months were recorded. The dietary data were evaluated by determining the consumed quantity and multiplying the portion size by the corresponding consumption frequency as reported in the FFQ. Standard portion sizes for women and men, separately, were used in this calculation; all food items in the FFQ were linked to food items in the Danish Food Composition Databank as previously described.[19]. Estimation of daily intake of macro-and micronutrients for each participant was based on calculations in the software program FoodCalc version1.3 (http://www.ibt.ku.dk/jesper/FoodCalc/Default.htm).

### Sample collection

Fecal sample collection and analysys was performed as previously described.[4] Urine was collected at the first visit upon arrival at the study site and stored at −80°C until analysis.

### Metabolic profiling

Urine samples were randomized, prepared and measured on a NMR spectrometer (Bruker) operating at 600.22 MHz ^1^H frequency using previously published experimental parameters[23] The ^1^H NMR spectra were then pre-processed and analyzed as previously reported[10] using Statistical Recoupling of Variables-algorithm.[24] Structural assignment was performed as reviewed in [25], using in-house and publicly available databases.

### Metagenomics

Shotgun sequencing of microbial DNA and metagenomics processing workflow for gene richness was performed as previously published.[4] Sequences were mapped onto the previously released integrated gene catalog.[2] Following the previously published strategy,[26] we built a novel set of manually curated gut-specific metabolic modules (GMMs) to describe and map microbial phenylpropanoid metabolism from shotgun metagenomic data. The set of 20 modules, following KEGG syntax, is provided in supplement, including references to the original publications where pathways were discovered and described (***Supplementary List***).

### Univariate statistical analysis

A ROUT test was performed to identify potential outliers (Q=1%). For statistical comparisons between study groups, normality was tested using D’Agostino-Pearson omnibus normality test, then one-way ANOVA was used, followed by Tukey’s HSD post hoc testing when data were normally distributed, otherwise groups were compared using the two-tailed Mann-Whitney test. Data are displayed as mean ± s.e.m in all figures. Multiple testing corrections were performed using Storey’s procedure.[27]

### Multivariate statistics

Probabilistic principal component analysis (PCA) was performed using MATLAB R2014a function ‘ppca’ to handle missing values. Unconstrained ordination was performed using principal coordinates analysis (PCoA) to visualize inter-individual variation in microbiota composition using Bray-Curtis dissimilarity on the genus-level abundance matrix using the R package *vegan*.[28] Distance based Redundancy Analysis (dbRDA) was used to determine the cumulative contributions of a matrix of explanatory variables on the response data, hippurate excretion inter-individual variation (Euclidian distance on log-transformed urine hippurate concentration matrix) in R package *vegan.*[28]. Orthogonal partial least squares discriminant analysis (O-PLS-DA) was performed in MATLAB R2014a for supervised multivariate analysis as previously described.[29] The predictive capability of O-PLS-DA models was evaluated through 7-fold cross-validation[29] to compute Q^2^_Yhat_ goodness-of-prediction parameters. The empirical significance of the Q^2^_Yhat_ parameter was evaluated by random permutation testing (10,000 iterations).[30]

### Animal experiments

Six-week-old C57BL/6J male mice (Janvier Labs, Courtaboeuf, France) were maintained in specific pathogen free condition on a 12h light/dark cycle and fed a standard chow diet (R04-40, Safe, Augy, France) for a week. Groups of 10 randomly selected mice were then fed either control chow diet (CHD) (D 12450Ki, Research diets, NJ) or high fat (60% fat and sucrose) diet (HFD) (D12492i, Research diets, NJ). One week later, osmotic minipumps (Alzet® model 2006, Charles River Lab France, l’Arbresle, France) filled with a solution of either hippuric acid or benzoic acid (5.55mM in 0.9% NaCl) (Sigma Aldrich, St Quentin, France) were inserted subcutaneously under isoflurane anesthesia. The metabolites were delivered at a rate of 0.15 µL/hour over a 42-day period (20 nmol/day).

Glycemia and body weight were recorded weekly. After 3 weeks of metabolite treatment, mice underwent an intra-peritoneal glucose tolerance test (IPGTT, 2g/kg). Blood was collected from the tail vein before glucose injection and 15, 30, 60 and 120 minutes afterwards to determine glycemia using an Accu-Check® Performa (Roche Diagnostics, Meylan, France). Cumulative glycemia (AUC) was calculated as the sum of plasma glucose values during the IPGTT and cumulative glucose increase (ΔG) parameter was calculated as AUC above the fasting glycemia baseline integrated over the 120 min of the IPGTT. Insulinemia was determined at baseline and at 30 minutes using insulin ELISA kits (Mercodia, Uppsala, Sweden). After 6 weeks of metabolite infusion, mice were killed by decapitation and organs were dissected and weighed. Triglycerides were quantified using a colorimetric assay (ab65336, Abcam, Paris, France) of liver homogenates. All procedures were authorized following review by the institutional ethic committee and carried out under national licence condition (Ref 00486.02).

### Histology and immunohistochemistry of animal tissues

Liver fibrogenesis and immunohistochemistry were determined as previously described [31]. An epitope-specific antibody (C27C9) was used for immunohistochemistry detection of insulin on pancreas sections (8508S Ozyme, Saint Quentin en Yvelines, France).

### RNA isolation and quantitative RT-PCR

Liver RNA preparation and reverse transcription were performed as previously reported [31]. Quantitative RT-PCRs were performed using oligonucleotides designed for the genes *Col1* (Forward: CACCCCAGCGAAGAACTCATA; Reverse: GCCACCATTGATAGTCTCTCCTAAC) and *Col3* (Forward: GCACAGCAGTCCAACGTAGA;

Reverse: TCTCCAAATGGGATCTCTGG) and using the cyclophilin A housekeeping gene.[31]

## RESULTS

### Hippurate is the urine metabolite most strongly associated with fecal microbial gene richness

To identify microbial and host compounds mediating beneficial effects in metabolic health, we profiled the urinary metabolome of the MetaHIT population[4] using ^1^H nuclear magnetic resonance (NMR) spectroscopy to perform a Metabolome-Wide Association Study (MWAS)[11] for microbial gene richness, a proposed criterion of metabolic and immune health[4] (Figure 1). We first built an orthogonal partial least squares (O-PLS) regression model based on the ^1^H NMR spectra to stratify the population by gene richness quartiles computed using our previously published 10-million integrated gene catalog (IGC)[2] (Figure 1A, P=5.8×10^-21^). The cross-validated model significantly predicted variance associated with gene richness through a permutation test (Figure 1B, P=9.7×10^-5^, 10,000 randomizations). Model coefficients for this discrimination revealed hippurate as having the strongest association with microbial gene count (Figure 1C): individuals with low microbial richness present significantly lower urinary hippurate levels than individuals with high microbial richness (Figure 1D, P=1.99×10^-9^, r^2^=0.173). These data support reports of association between hippurate levels and microbial functional redundancy[26] and Shannon’s diversity index[17] (Figure 1E, P=0.024 and Supplementary Figure 1A, P=0.0058).

**Figure 1.**
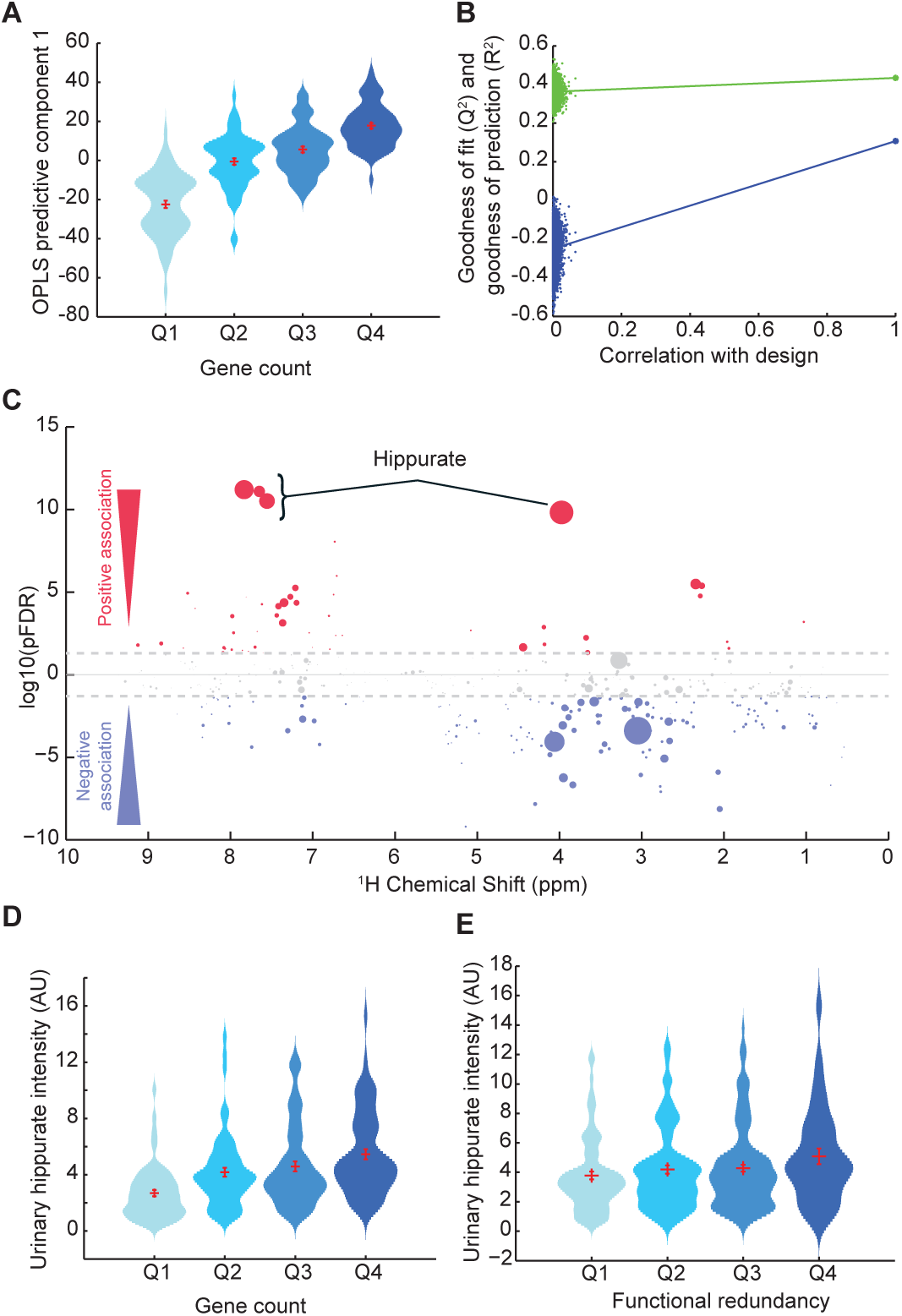
Hippurate is the main metabolite correlated with gene richness and functional redundancy of the microbiome. **(A)** Scores plot (predictive component 1) obtained for an O-PLS-DA model fitted usingurinary^1^H NMR-spectra to predict microbial gene richness, showing a significant association between gene richness quartiles and ^1^H NMR spectra (p=5.84×10^-21^ for a significantly non-zero slope using F-test, n=271). **(B)** Empirical assessment of the significance of O-PLS goodness-of-fit parameter Q^2^_Yhat_ by generating a null distribution with 10,000 random permutations (p=9.68×10^-5^). **(C)** Manhattan plot highlighting associations between ^1^H NMR variables and gene count displayed in a pseudo-spectrum layout. A negative value (blue circles) means a negative correlation while a positive value (red circles) means a positive correlation. Grey circles are clusters with a p-value>0.01. Size of circles represents the covariance of the cluster with the gene count.**( D)** Association between urinary hippurate intensity and gene count quartiles (p=1.99×10^-9^ for a significantly non-zero slope using F-test). **(E)** Association between urinary hippurate intensity and microbial functional redundancy [26] quartiles (p=0.0239 for a significantly non-zero slope using F-test, n=271).

Consistent with associations previously reported for microbial gene richness in the MetaHIT population[4] and associations between hippurate and reduced cardiometabolic disease risk[11,12,14,17], urinary hippurate significantly correlated with low values for body mass index (BMI), bodyweight, the homeostasis model assessment of insulin resistance (HOMA-IR) and fasting circulating levels of IL6, insulin, and C-peptide, whilst adjusting for age and gender (partial Spearman’s rank-based correlations, q<0.1, Supplementary Figure 1B). Moreover, stratification on urinary hippurate concentrations in lean (BMI <25 kg/m^2^), and overweight or obese (BMI >25 kg/m^2^) individuals showed improved glucose homeostasis in participants excreting higher levels of hippurate (Supplementary Figure 1C, *median threshold*). These observations depict hippurate, one of the main microbial-mammalian co-metabolites found in human urine, as a key molecular marker associated with gene richness which may underlie some of its health benefits. These results however raise questions related to the entanglement of gene richness and hippurate as potential markers of health[13, 17]. Adjusting for hippurate weakens associations between gene richness and bioclinical variables (Supplementary Figure1B), consistent with the idea that hippurate could mediate some of the observed benefits for subjects with higher gene richness. However, hippurate associations with bioclinical variables adjusted for gene richness are no longer significant, suggesting that the gene richness signal overrides hippurate associations in the presence of confounding variation affecting urinary concentrations, such as diet, microbial synthesis, host conjugation and clearance. Hippurate did not correlate either to glycine bioavailability, which is required for hippurate synthesis through conjugation with gut microbial benzoate[32] or kidney function (eGFR) which could limit hippurate synthesis and clearance (Supplementary Figure 1D-E).

### Microbiome determinants of hippurate production in the phenylpropanoid pathway

To characterize the microbial determinants of benzoate production, we next focussed on high-throughput shotgun sequencing fecal metagenomics data (n=271). We functionally annotated functions of the IGC to KEGG Orthology (KO) groups and found 2,733 KEGG and 6,931 EggNOG modules positively associated with urine hippurate levels (pFDR<0.05, Supplementary Tables S1-2). Of specifically curated enzymatic modules[26] involved in microbial benzoate metabolism (4 aerobic and 15 anaerobic; Supplementary List), only three modules significantly correlated with urine hippurate levels: MC0004 (detected in 271 samples) corresponding to a 2-enoate reductase converting cinnamic acid into 3-(3-hydroxyphenyl)-propionic acid, MC0005 (observed in 201 samples) converting cinnamate into benzoate and MC0014, a benzoate 4-monooxygenase only observed in fewer than 15% of individuals (Figure 2A, Supplementary Table S3). Abundance of these modules also correlated with gene richness (Supplementary Table S4), thereby providing a functional basis for the association between gene richness and urine hippurate levels observed in this population (Figure 1). Genes involved in MC0004 and MC0005 were predominantly found in genomes from unclassified Firmicutes and Clostridiales (Figure 2B, Supplementary Tables S5-6). Among classified Firmicutes, the genera *Lachnoclostridium*, *Eubacterium* and *Blautia* harbored MC0004. Conversely, MC0005 was encoded by Proteobacteria, i.e. *Klebsiella*, *Enterobacter*, *Suterella* and *Comamonas* (Figure 2B). We then mapped these modules into the enteroscape (as observed on the principal coordinates plot derived from normalized genus abundances using Bray-Curtis distances,[26] Figure 2C), revealing that the conversion of cinnamic acid into 3-hydroxy-3-phenylpropionic acid is linked to the *Ruminococcus* enterotype, while capacity to convert cinnamate to benzoate is more ubiquitously distributed across gut community types. Phenylpropanoid pathway potential is increased in the *Ruminococcaceae*-enterotype compared to the *Bacteroides*- or *Prevotella*-enterotypes, the former being confirmed by analyzing gradients of key taxa instead of enterotypes (Supplementary Table S7). The results altogether suggest a wide range of substrates, taxa and species are involved in benzoate production, consistent with its association with gene richness.

**Figure 2.**
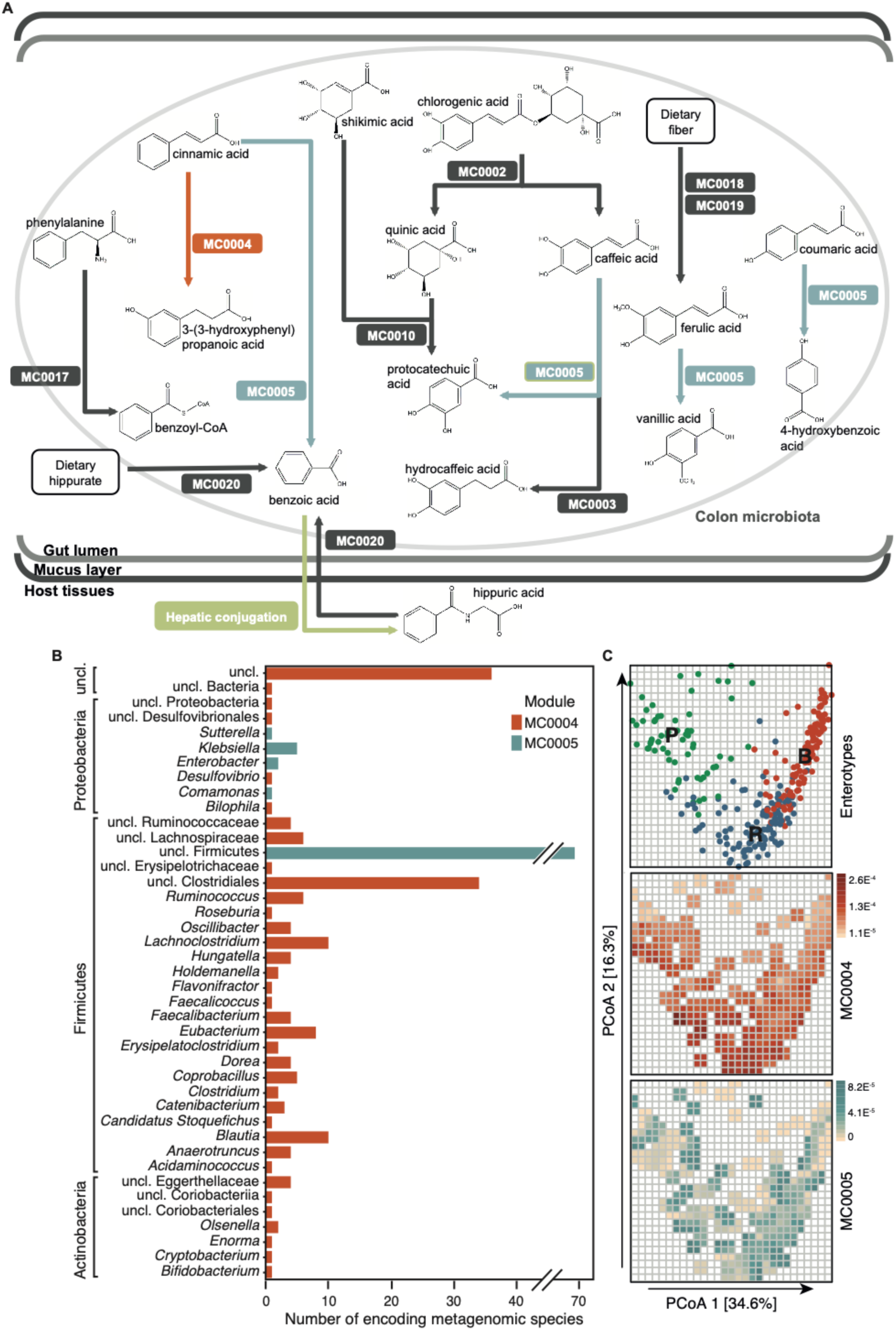
Detection of microbial phenylpropanoid metabolism-related modules in fecal metagenomes of healthy volunteers and their associations with urine hippurate concentrations. **(A)** Visualisation of gut-specific metabolic modules (GMMs) encoding anaerobic phenylpropanoid metabolism-related pathways detected in more than 20% of individuals; MC0004 (orange; Spearman rho=0.19, q-value=0.005) and MC0005 (blue; Spearman rho=0.21, q-value=0.005) correlate positively to urine hippurate concentrations (n=271). All metabolites are connected to benzoate but for clarity the non-significant reactions were omitted. **(B)** Metagenomic species encoding modules MC0004 and MC0005. **(C)** [top panel] Fecal microbiomes dissimilarity visualised on the first plane of the genus-level principal coordinates analysis (PCoA, Bray-Curtis dissimilarity), with individual samples colored according to enterotypes (R, Ruminococcaceae; B, Bacteroides; P, Prevotella). [middle and bottom panels] Same genus-level PCoA overlaid with a mesh colored according to the median abundances of GMMs MC0004 (red) and MC0005 (blue) in samples falling within each cell of the mesh (n=271). See Supplementary Table S3 for correlation between hippurate and GMMs.

### Urine hippurate levels associate with improved metabolic health in patients with diets rich in meat and saturated fats

We next assessed individual nutritional intake through validated FFQs available in 193 MetaHIT individuals.[19] Associations with metabolic health were stratified according to multivariate dietary patterns (Figure 3). A PCA of 133 dietary intake descriptors summarises dietary patterns and loadings highlight four archetypal diets within our population: higher consumption of fruits and vegetables (n=96) vs. high consumption of meat containing saturated fats (n=97) on the first principal component (PC1) and carbohydrate-rich foods vs. fish containing unsaturated fats on PC2 (Figure 3A), a trend which was observed at the food ingredient and nutrient level (Supplementary Figure 2A-B). It is therefore possible to stratify the population according to the median of dietary PC1 highlighting contrasts between healthy (higher consumption of fruit and vegetables) and at-risk (higher consumption of carbohydrates and meat) diets. Although hippurate was not correlated with the first two dietary principal components, its dynamic range was similar whilst stratifiying according to the first two principal components (Supplementary Figure 2C-E). To summarise the main factors influencing inter-individual variation in urine hippurate excretion, we calculated the cumulative contribution of several covariates using a dbRDA ordination approach (Figure 3B). Gene richness accounted for 12% (P=0.002), followed by MC0020 encoding a hippurate dehydrolase (4%, P=0.002, observed in 271 subjects) catalysing the retroconversion of hippurate into benzoate, and HOMA-IR (1.5%, P=0.008; n=265). When taking diet into account (i.e., PC1 fruits and vegetables vs. meat; n=193) in the dbRDA, gene richness contributes to 15% (P=0.002), diet adding another 4% (P=0.002) and hippurate retroconversion 3% (P=0.004), suggesting that the pattern of hippurate associations could be diet-dependent and requiring further analysis. We therefore stratified the data according to diet (n=193) using a median threshold for the first dietary principal component, highlighting a healthy diet associated with vegetables and fruit intake (low PC1 values, n=96) and an at-risk diet rich in saturated fats derived from meat intake (n=97). For this latter subset of individuals consuming a diet rich in saturated fats on the first dietary principal component, urine hippurate levels significantly associated with decreased HOMA-IR, circulating fasting levels of insulin, fasting associated adipocyte factor (FIAF, also known as Angiopoietin-like 4, a peroxisome proliferator-activated receptor target gene environmentally modulated by the gut microbiota inhibiting lipoprotein lipase in adipose tissue [33]) and TNF-*α*, whilst displaying increased plasma levels of fasting adiponectin (Figure 3C-E, Supplementary Table S8). Urine hippurate was not associated with any health benefits in the subsets of participants consuming mostly a fruit and vegetable diet, a pescetarian diet or a carbohydrate-rich diet (Supplementary Table S8).

**Figure 3.**
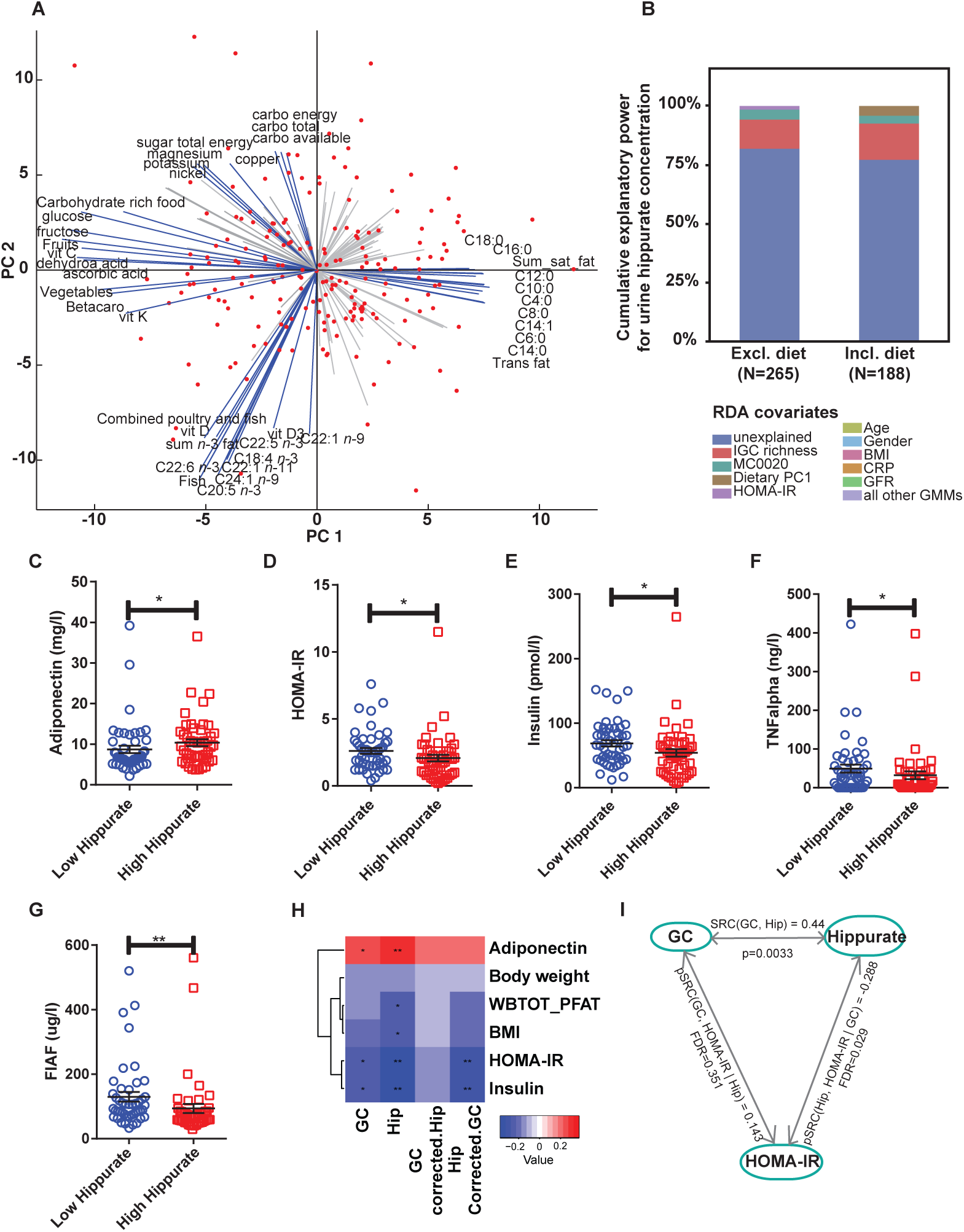
Hippurate associates with improved glucose homeostasis only in participants consuming a diet rich in saturated fats. **(A)** Biplot of the principal component analysis (PCA) of dietary intakes highlights opposite diets along first two principal components (PCs). The main drivers of each principal components are named and represented by blue arrows. **(B)** Cumulative contributions of explanatory variables to inter-individual variation in hippurate excretion, estimated by redundancy analysis (dbRDA). Explanatory variables included microbiota gene count, microbiota phenylpropanoid metabolism modules, host dietary principal components and clinical parameters (age, gender, BMI, HOMA-IR, CRP, serum glycine levels, and glomerular filtration rate (eGFR) estimation with CKD-EPI). **(C-F)** Evaluation of hippurate stratification (high hippurate, n=49 vs low hippurate, n=48) on bioclinical variables (q<0.1, Supplementary Table S8) for individuals on high PC1 (i.e. high meat / high saturated fat diet). For full name description of physiological data see Supplementary Table S8.

To disentangle contributions arising from hippurate and gene richness to bioclinical variables in subjects consuming a diet rich in fats, we compared unadjusted and adjusted Spearman’s rank-based correlations (Figure 3H). In the population consuming higher amounts of meat and saturated fats, elevated urine hippurate levels significantly associated with an increase in fasting plasma adiponectin and a reduction in adiposity, BMI, HOMA-IR and fasting plasma insulin, which is consistent with gene richness being significantly associated with an increase in adiponectin and a decrease in HOMA-IR and fasting plasma insulin. However, the associations between gene richness and bioclinical variables were no longer significant when adjusting for urine hippurate levels. Conversely, hippurate associations with insulin and HOMA-IR were still significant after gene richness adjustment. We exemplified this through a correlation graph taking into account the correlation between hippurate and gene richness (r=0.44): this unadjusted correlation between gene richness and HOMA-IR collapses when adjusting for gene richness (rho=0.143, n.s.) and is in fact contributed for by a partial correlation between urine hippurate and HOMA-IR (Figure 3I). The latter finding suggests a possible preventive role for hippurate in metabolic disease driven by diets high in meat and saturated fats, independently of gene richness.

### Hippurate and benzoate improve glucose tolerance in HFD-fed mice

To further study the impact of benzoate and hippurate on host physiology, we treated mice with subcutaneous infusion of hippurate (0.14 mg/kg/day) and benzoate (0.1 mg/kg/day) in CHD and HFD (Figure 4). Under control diet, metabolite treatments had no effect on body weight, BMI and fasting glycemia (Supplementary Figure 4). Benzoate caused a significant elevation of the adiposity index and a reduction of the normalised heart weight (Supplementary Figure 5A). During an IPGTT, both metabolites induced a slight increase in glycemia (Figure 4A-B) and ΔG parameter (Figure 4C), respectively. Also, hippurate induced a stronger glucose-stimulated insulin secretion than benzoate, compared to the saline-treated mice (Figure 4D). Whilst HFD-feeding increased body weight and adiposity, hyperglycemia and glucose intolerance (Figure 4E-H, Supplementary Figs 4D,E and 5), mice treated with hippurate or benzoate showed a parallel improvement in glucose tolerance compared to saline-treated controls (Figure 4E). This effect was illustrated by the highly significant reduction of both the cumulative glycemia during the test (Hippurate vs. control −23.90% P=0,001, benzoate vs. ctl −31.52%, P=0.001) and the ΔG parameter (Hippurate vs. ctrl −37.22% P=0,001, benzoate vs ctrl −33.35%, P=0.001) (Figure 4F,G). Hippurate and benzoate treatments also significantly increased glucose-induced insulin secretion (Figure 4H). These data suggest that both metabolites have the capacity to improve glucose tolerance and stimulate glucose-induced insulin secretion *in vivo* specifically in diet-induced obesity and diabetes.

**Figure 4.**
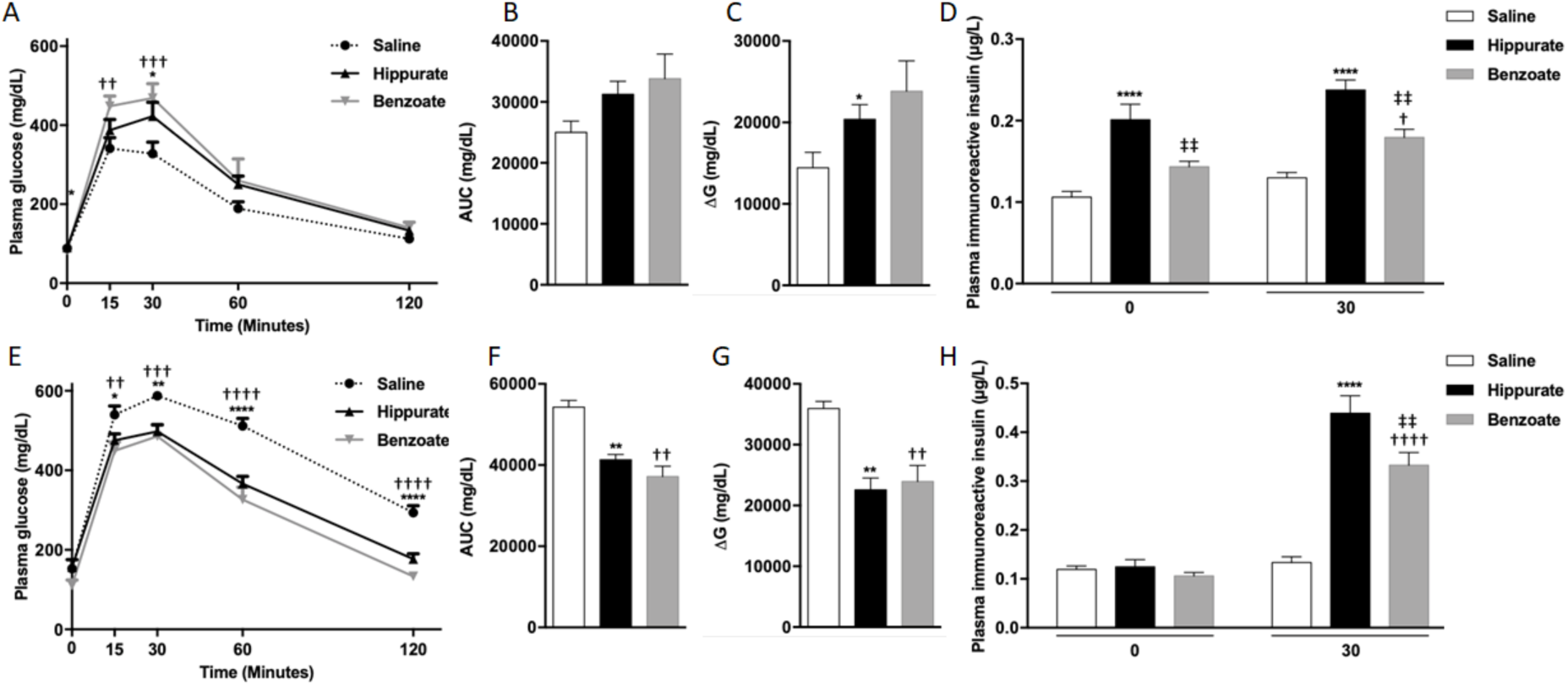
Chronic hippurate treatment improves glucose homeostasis in HFD-fed mice. **(A)** Heatmap summarising Spearman’s partial correlation between gene richness, hippurate,gene richness adjusted for hippurateand hippurate adjusted for gene richness and bioclinical variables, all correlations adjusted for age and gender. Stars represent significant pFDR corrected using Benjamini Hochberg procedure * pFDR<0.1, ** pFDR<0.05, *** p<0.01. **(B)** Representation of the Spearman’s correlations and partial correlations between gene count and hippurate, hippurate and HOMA-IR adjusted for gene richness and between gene richness and HOMA-IR adjusted for hippurate. **(C)** Plasma glucose during a glucose tolerance test (GTT). **(D)** Area under the curve for glucose during the GTT. **(E)** Body weight of mice during the 6 weeks of hippurate treatment. **(F)** Body mass index at sacrifice **(G)** Adipose tissue weight normalised to body weight. **(H)** Plasma adiponectin concentration. For chow diet and chow diet + hippurate, (n=10) and for HFD and HFD + hippurate (n=6). Data shown are mean±SEM. Statistical analysis was performed using two-way ANOVA with Tukey’s posthoc test. ** p<0.01, **** p<0.0001. For panel (D), (F), (G) and (H), groups with different superscript letters are significantly different (P<0.05), according to Tukey’s posthoc test.

**Figure 5.**
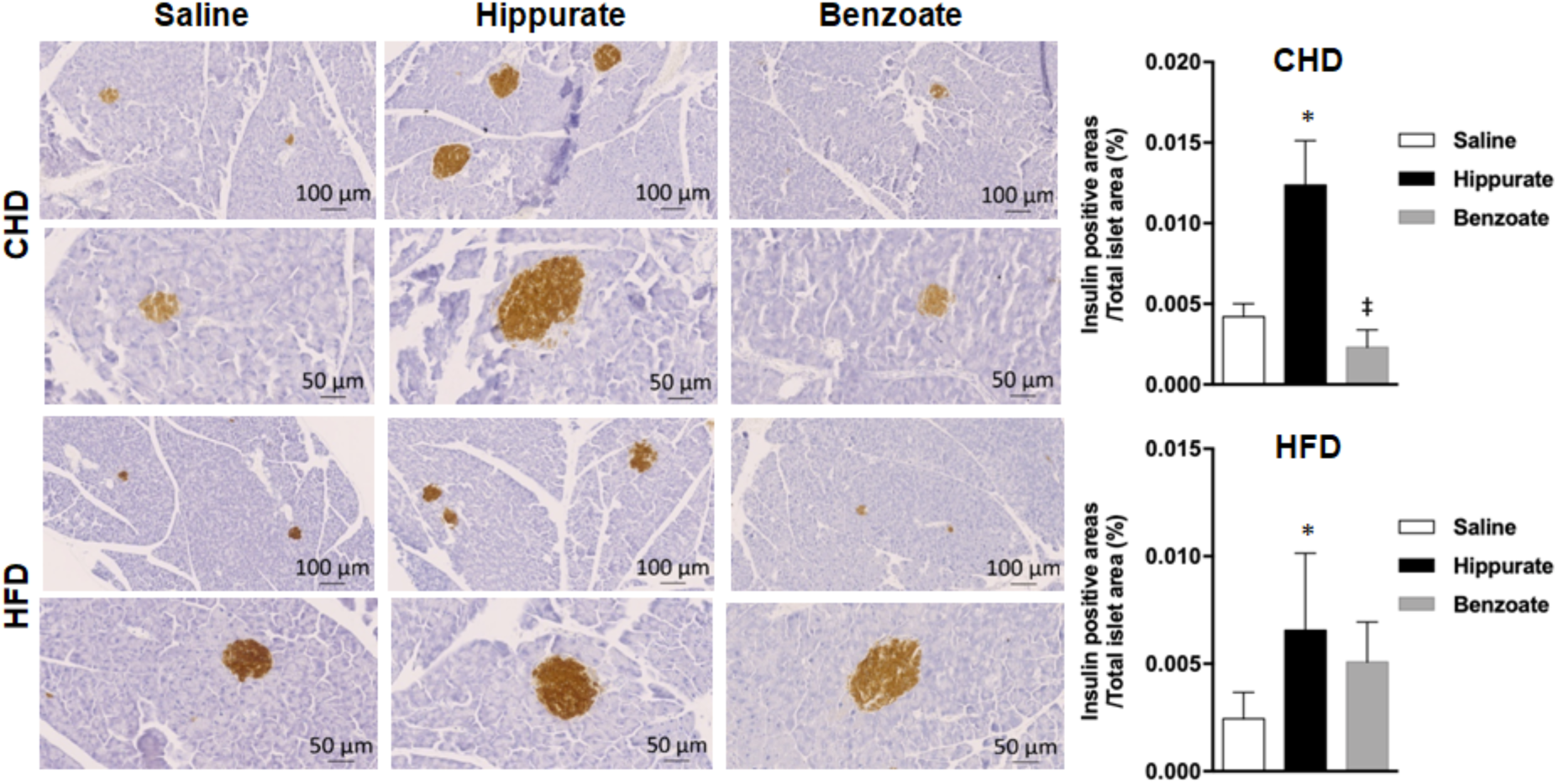
Effect of chronic administration of hippurate and benzoate on pancreatic islets in C57BL6/J mice. The effect of chronic subcutaneous administration of the metabolites (5.55 mM) for 42 days on islet density was tested in mice fed chow diet (CHD) or high fat diet (HFD) for 56 days. Control mice were treated with saline. Each biological replicate represents one slide per animal mounted with at least 3 tissue sections, representing 3 technical replicates, the mean and variance of which is presented as the result per biological replicate. Results are expressed as percentage of positive pixels. ‡P<0.05, significantly different between mice treated with benzoate and hippurate.

### Effects of hippurate and benzoate on the morphology of pancreatic islets

To investigate the possible cause of stimulated insulin secretion by hippurate and benzoate, we performed out a histological analysis of pancreatic islet structure. We confirmed that insulin positive area was increased by hippurate (+194%, P=0.04, one-tailed) or benzoate (+437%, p=0.02, one-tailed) respectively in mice fed control diet (Figure 5); hippurate treatment Insulin positive area was also increased in HFD-fed mice treated with hippurate (+168%, P=0.04, one-tailed). These data suggest that increased β-cell mass may explain enhanced insulin production and secretion induced by hippurate and benzoate treatment.

### Effects of hippurate and benzoate on liver histopathology and function

Liver triglyceride accumulation, fibrosis and inflammation are hallmarks of structural and biochemical adaptations to HFD feeding. Liver triglycerides were more elevated in HFD fed mice (29.30±4.15 mg/g) than in mice fed control diet (8.63±2.19 mg/g, P=0.002) (Supplementary Figure 6). Hepatic triglycerides were not significantly affected by hippurate or benzoate treatment in either diets. Benzoate induced a singifnicant decrease in liver triglycerides compared to hippurate in HFD (−57.35%, P= 0.02) (Supplementary Figure 6). We next analysed hepatic fibrosis through quantitative analysis of collagen detected by red picrosirius staining of histological sections (Figure 6A). Hippurate treatment resulted in a marked reduction of liver collagen in mice fed control diet (−53.2%) or HFD (−55.7%), whereas benzoate had no effects on collagen levels in these mice (Figure 6B,C). These results were confirmed by liver expression of the genes encoding collagen 1 (*Col1*) and collagen 3 (*Col3*) (Figure 6B,C): hippurate treatment induced a significant reduction of the expression of *Col1* (−79.89%, P=0.02) and *Col3* (−29.37%, P=0.01) under control diet (Figure 6B). but the effect of the metaolites on *Col1* and *Col3* I HFD was marginal (Figure 6C). Finally, we assessed the effects of hippurate and benzoate on liver inflammation through *α*-SMA (alpha Smooth Muscle Actin) staining[34, 35] (Figure 7A), which is increased by HFD-feeding (+273.86%) (Figure 7B,C). Hippurate induced a marked reduction in *α*-SMA staining in mice fed control diet (−82.71%) or HFD (−94.87%) (Figure 7B,C), suggesting reduced presence of stellar cells and reduced liver inflammation when compared to saline-treated controls. In contrast, benzoate treatment in chow diet induced a strongly significant increase in stellar cells when compared to mice treated with saline (+564%, P=10^-7^) or hippurate (+3,741%, P=10^-7^), thereby indicating liver inflammation in these mice (Figure 7B). Collectively, these data show that hippurate decreases fibrosis and inflammation regardless of diet, whereas benzoate reduces triglycerides and collagen accumulation in obese mice fed HFD whilst stimulating inflammation in lean mice fed control diet.

**Figure 6.**
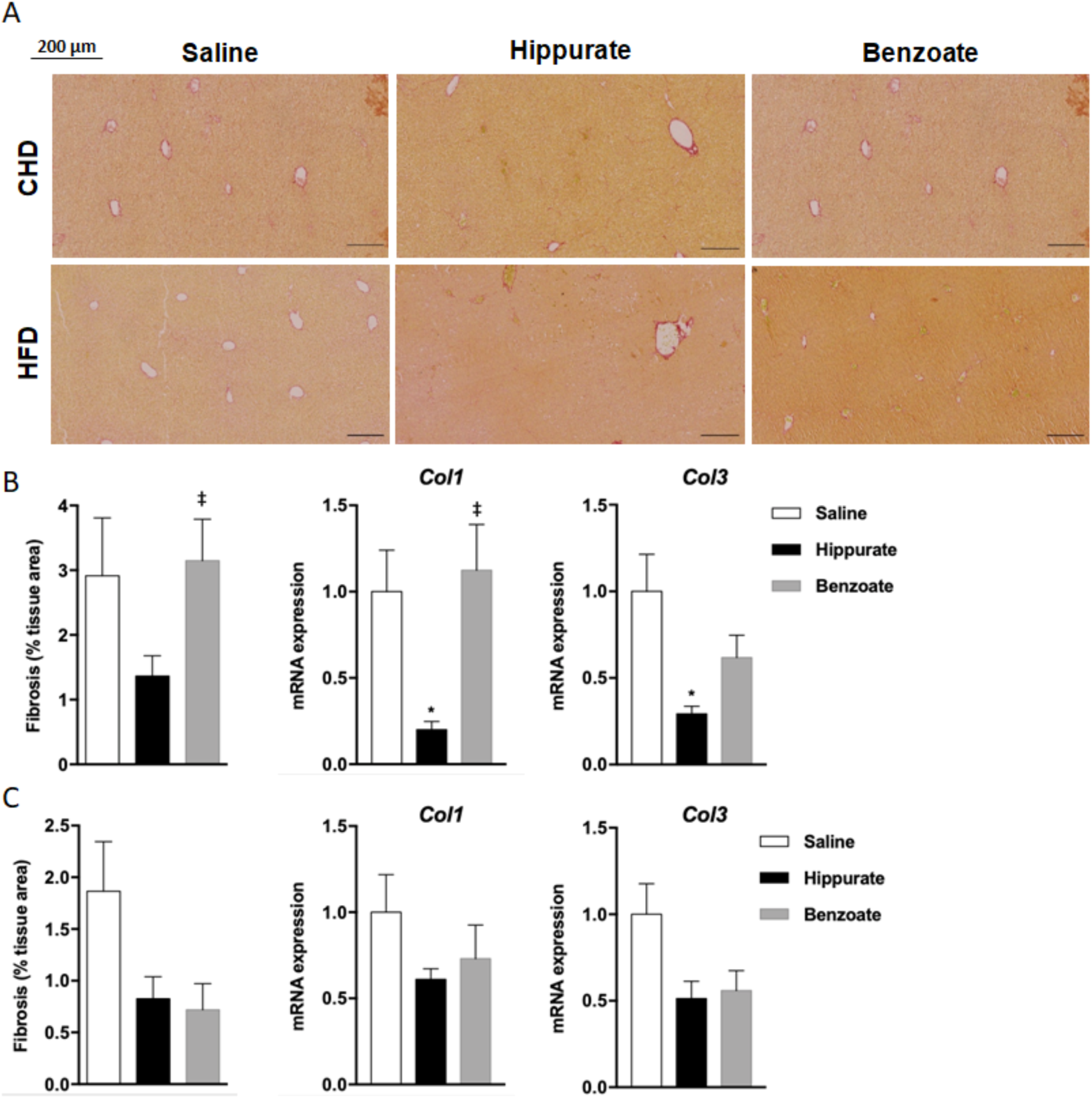
Effect of chronic administration of hippurate and benzoate on liver fibrosis in C57BL6/J mice. **(A)** The effect of chronic subcutaneous administration of the metabolites (5.55 mM) for 42 days on liver collagen was tested in mice fed control chow diet (CHD) or high fat diet (HFD) for 56 days. Control mice were treated with saline. Red Picrosirius staining of histological sections was used to visualise and quantify fibrosis in mice fed CHD **(B)** or HFD **(C)**. Each biological replicate represents one slide per animal mounted with at least 3 tissue sections, representing 3 technical replicates, the mean and variance of which is presented as the result per biological replicate **(B,C)**. Liver expression of the genes encoding collagen 1 (*Col1*) and collagen 3 (*Col3*) was assessed in mice fed CHD (B) or HFD **(C)** by quantitative RT PCR in 6 mice per group. Data were analyzed using the unpaired Mann-Whitney test. Results are means ± SEM. *P<0.05 significantly different between mice treated with hippurate and controls. ‡P<0.05, significantly different between mice treated with benzoate and hippurate.

**Figure 7.**
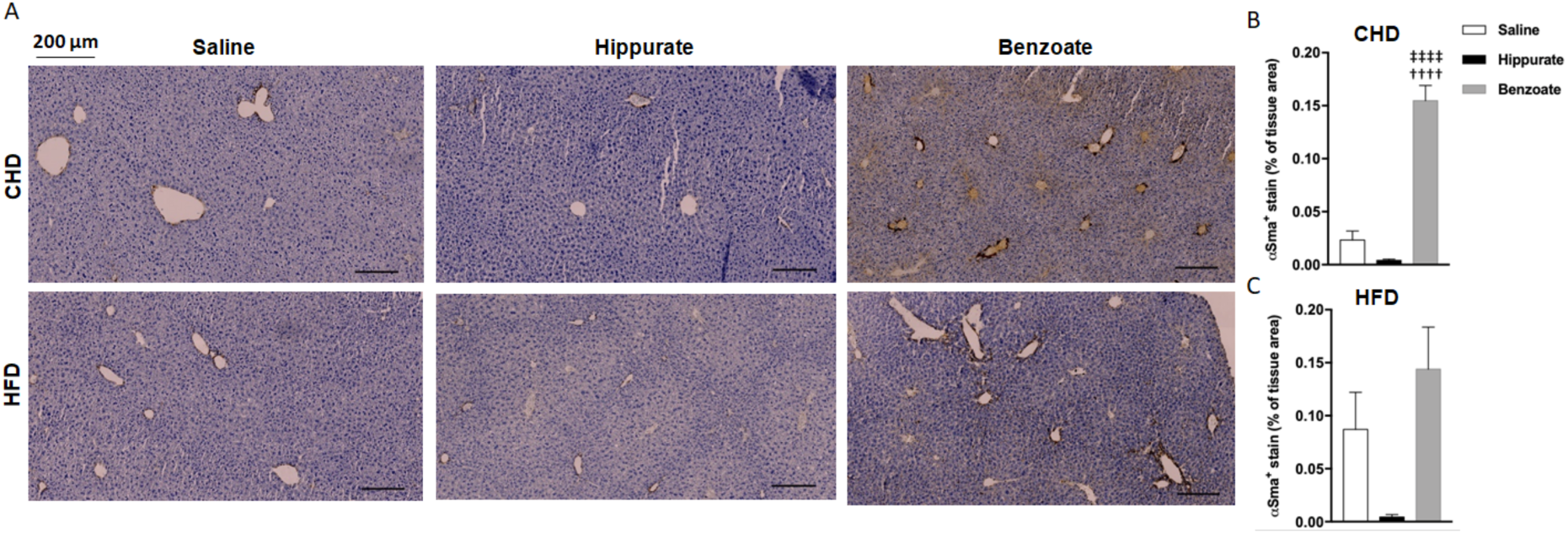
Effect of chronic administration of hippurate and benzoate on liver inflammation in C57BL6/J mice. αSMA staining of liver slides was used to assess inflammation in mice fed chow diet (CHD) or high fat diet (HFD) for 56 days and chronically treated with subcutaneous administration of the metabolites (5.55mM) for 42 days **(A)**. Control mice were treated with saline. Each biological replicate represents one slide per animal mounted with at least 3 tissue sections, representing 3 technical replicates, the mean and variance of which is presented as the result per biological replicate in mice fed CHD **(B)** or HFD **(C)**. Data were analyzed using the unpaired Mann-Whitney test. Results are means ± SEM. ††††P<0.0001, significantly different between mice treated with benzoate and saline treated controls. ‡‡‡‡P<0.0001, significantly different between mice treated with benzoate and hippurate.

## DISCUSSION

Integrated analysis of metabolome profiling and deep metagenome sequencing data of 271 middle-aged non-diabetic Danish subjects from the MetaHIT study sample [4] identified urinary hippurate as the metabolite most significantly associated with microbial gene richness based on the microbial IGC.[2] This observation largely confirms previous reports associating hippurate with gene richness in steatosis and bariatric surgery contexts[14, 36] as well as with increased gut microbial diversity as assessed by sequencing the 16S rRNA gene amplicon.[17] Hippurate having previously been inversely correated with blood pressure, obesity and steatosis,[11–14] this study highlights diet-dependent relationships between microbiota-host co-metabolism of benzoate and hippurate and health benefits associated with gene richness. Our in-depth dissection of the metagenomic determinants of urinary hippurate highlighted a series of richness-responsive gene modules functionally related to benzoate synthesis. These modules are shared across several enterotypes and taxonomic gradients. Population stratification analyses demonstrated that hippurate only benefits individuals consuming a diet rich in saturated fats. This hypothesis of a diet-dependent beneficial health effect of benzoate metabolism was confirmed *in vivo* in a preclinical model of HFD-induced obesity.

### Hippurate brings diet-dependent metabolic improvements in pancreas and liver

Our study shows that chronic hippurate treatment in a model of obesity induced by HFD-feeding reduces glucose intolerance, stimulates insulin secretion, enhances β-cell mass and reduces hepatic inflammation and fibrosis. Metabolomic studies have consistently shown inverse associations variations between hippurate levels and pathophysiological elements of the metabolic syndrome. Urinary hippurate is reduced in mouse models of insulin resistance[10] and in rat models of spontaneously occurring hypertension (SHR), type 2 diabetes (Goto Kakizaki, GK) and obesity (Zucker) or the WKY rat.[37–39] This is consistent with our previous reports showing an inverse association among hippurate, insulin resistance, hypertension, obesity or liver steatosis[10-12,14] and observations that hippurate exerts protective effects in β-cells.[40] We also showed in HFD-fed isogenic mice that urinary hippurate measured before a 3-week HFD induction predicts the future development of obesity, suggesting that functional variations in microbiome predicts disease risk independently of genetics.[41] Whilst both hippurate and benzoate have similar *in vivo* effects, including greatly improved glucose tolerance and stimulation of insulin secretion, only hippurate results in beneficial effects on increased β-cell mass or reduced liver fibrosis.

### The phenylpropanoid-benzoate-hippurate pathway in metabolic health

The range of responses observed in the animal model treated with hippurate and benzoate depict subtle and context-dependent microbiome–host interactions. Benzoate and its co-metabolite hippurate are the endproducts of several converging microbial biosynthetic pathways.[15] The phenylpropanoid pathway is a broad network of reactions connecting a wide range of dietary substrates such as phenylalanine, quinic acid, shikimic acid or chlorogenic acid for instance to 4-coumaryl-coA. These pathways lead to much simpler molecules, benzoate being their common endpoint. Dietary and microbial intermediates (including cinnamic acids, coumarins, stillbenoids, flavonoids and isoflavonoids) in the phenylpropanoid and connected pathways are associated with beneficial health outcomes.[15, 42]

### Conclusion

Our study depicts hippurate as a pivotal microbial-host co-metabolite mediating part of the beneficial metabolic improvements associated with high microbial gene richness in the context of Western-style diets. This work unifies previous reports in which hippurate was associated improvements in insulin resistance, steatosis, hypertension and obesity[10-12,14] and microbial ecological diversity.[14, 17] Beyond the diversity of microbial ecosystems and functions associated with hippurate, we uncover beneficial bioactivities in the liver and pancreas resulting in health benefits in terms of metabolic control, liver inflammation and fibrosis. Our observations support the existence of diet-dependent microbial-host metabolic axis in which hippurate partly offsets unhealthy diets, further exemplifying that the microbiome determines key components of human biochemical individuality[43] and provides critical diagnostic and therapeutic potential in personalized nutrition and stratified medicine.[44, 45]

## Acknowledgements

This research was funded by FP7 METACARDIS HEALTH-F4-2012-305312 with additional funding from the Metagenopolis grant ANR-11-DPBS-0001 and support of the NIHR Imperial Biomedical Research Centre. The Novo Nordisk Foundation Center for Basic Metabolic Research is an independent research institution at the University of Copenhagen partially funded by an unrestricted donation from the Novo Nordisk Foundation. S.V.S. is funded by Marie Curie Actions FP7 People COFUND Proposal 267139 (acronym OMICS@VIB) and the Fund for Scientific Research-Flanders (FWO-V). L.H. was an MRC Intermediate Research Fellow (MR/L01632X/1).

## Contributions

F.B. J.C., T.N. and S.V.S. and contributed equally to this work. F.B. J.C. and G.I.M. acquired data, J.C., T.N., S.V.S. performed analyses, G.F. and S.V.S. curated the gut-specific metabolic modules, L.H., A.L.N., A.R.M., N.P., S.F., E.L.C., and A.M.L.L. participated in data collection and processing, J.C., T.N., S.V.S., F.B. and G.F. performed statistical analyses. M.-E.D., D.G., J.R. and O.P. designed the study. T.H., J.K.N. K.C., P.B., S.D.E., participated in the study design and interpretation of the results, M.E.D. wrote the manuscript with contributions from F.B., J.C., T.N., S.V.S, G.F., J.R., O.P and D.G.

## SUPPLEMENTARY SECTION

**Supplementary Figure 1.**
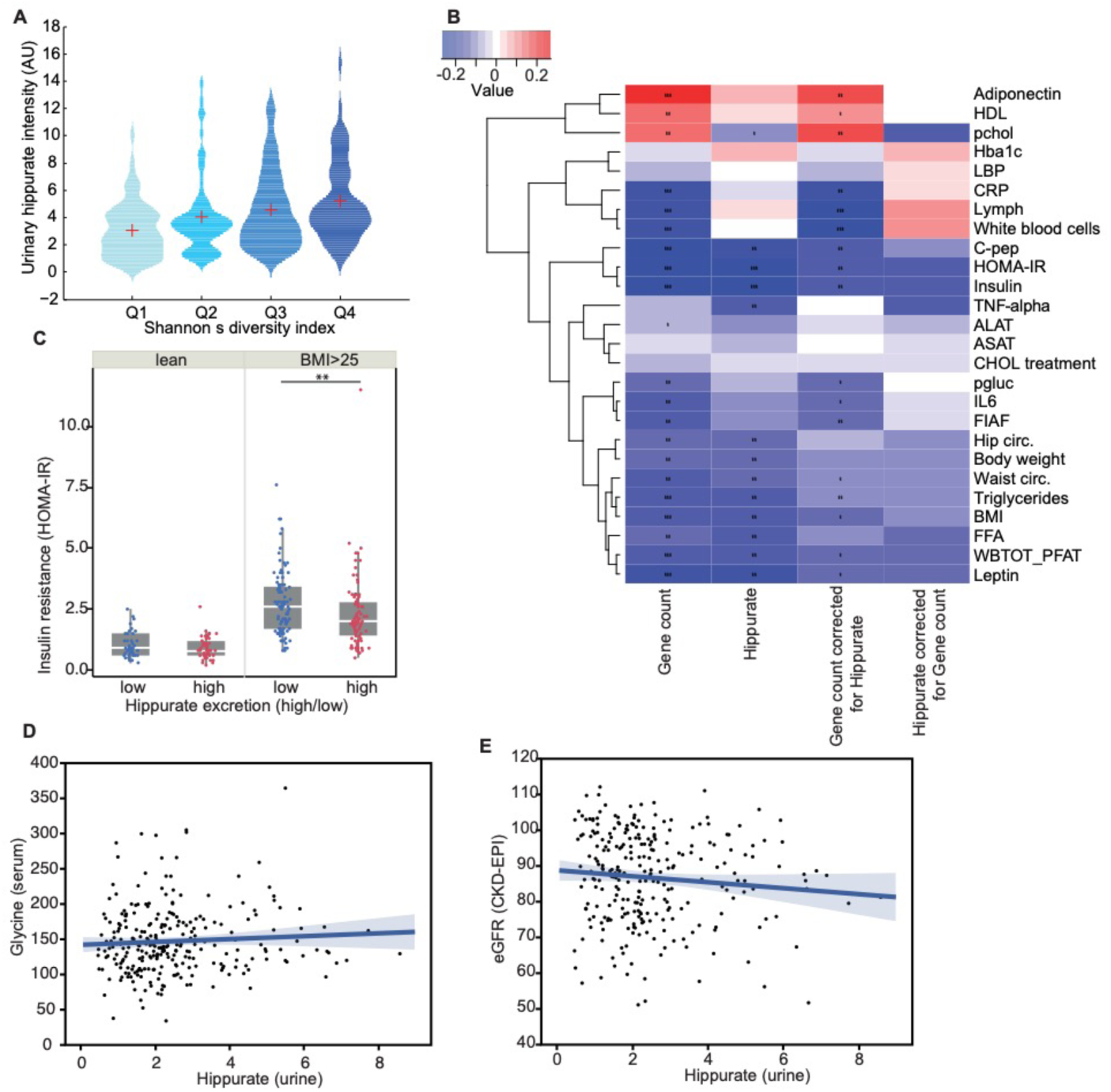
Relationship between gene richness, hippurate and bioclinical variables. **(A)** Association between urinary hippurate intensity and Shannon’s diversity index quartiles (p=6.04×10^-8^for a significantly non-zero slope using F-test, n=271). **(B)** Heatmap summarising significant (pFDR<0.1) Spearman’s correlation FDR corrected using Storey’s procedure[27] betweengene count, hippurate and gene count adjusted for hippurate and physiological data, all adjusted for age and gender.For full physiological data names and units see *Supplementary Table 8*. **(C)** Association between hippurate and insulin resistance (HOMA-IR) stratified according to BMI (lean (BMI≤25, n=87), overweight and obese (BMI>25, n=184) and hippurate excretion levels (Mann-Whitney U test, p=0.0058). **(D)** Representation of the absence of significant correlation between urinary hippurate and eGFR. **(E)** Representation of the absence of significant correlation between urinary hippurate and circulating glycine.

**Supplementary Figure 2.**
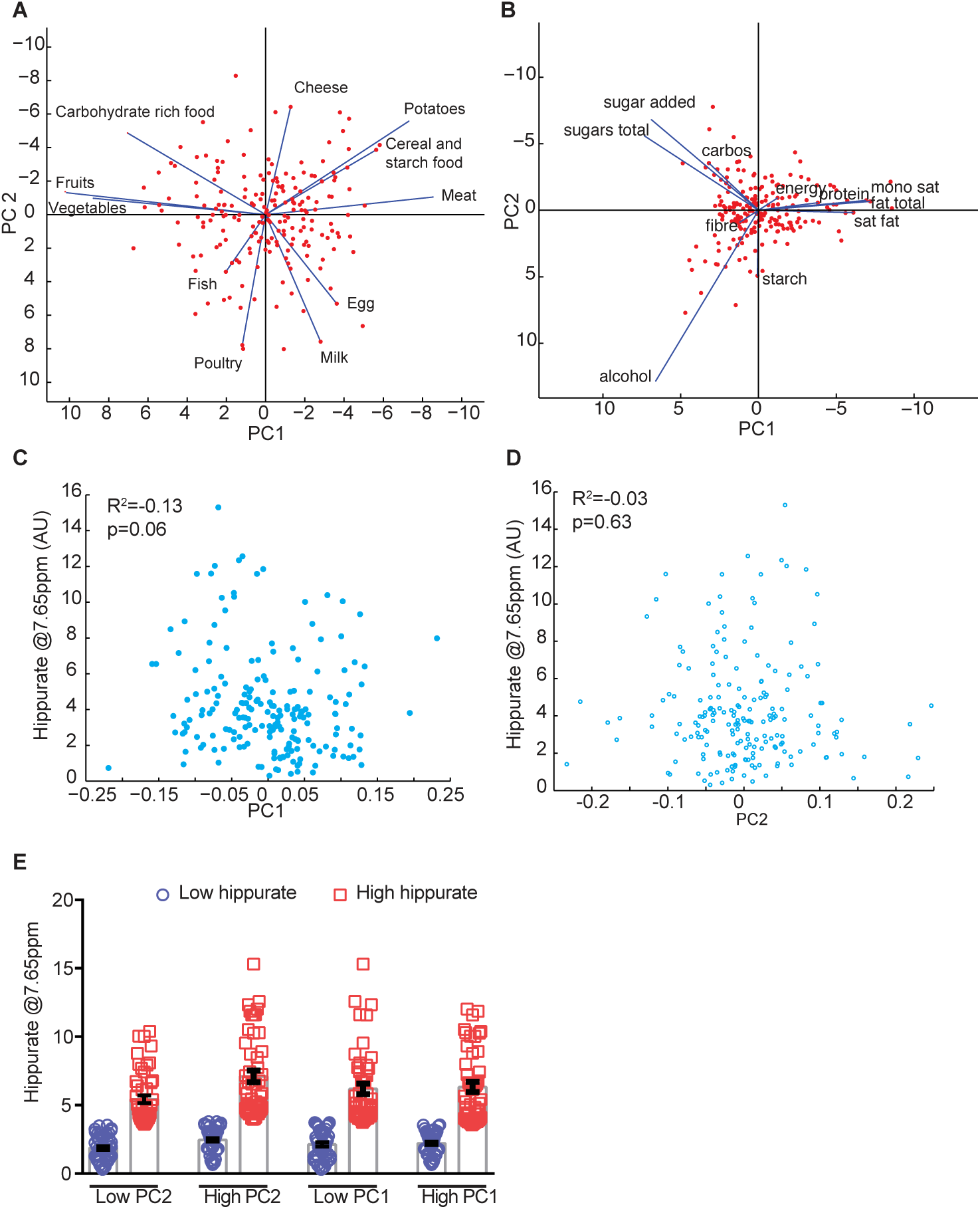
Urinary hippurate does not correlate with main dietary trends summarized and presents a high variability within each subgroup. **(A)** Biplot of the dietary data generated using only food items. **(B)** Biplot of thmicrobial e dietary data generated using only macronutrients items. **(C)** Representation of the absence of correlation between hippurate concentration and principal component 1. **(D)** Representation of the absence of correlation between hippurate concentration and principal component 2. **(E)** Distribution of hippurate urinary concentration within each individualsubgroup stratified in high and low hippurate using median.

**Supplementary Figure 3.**
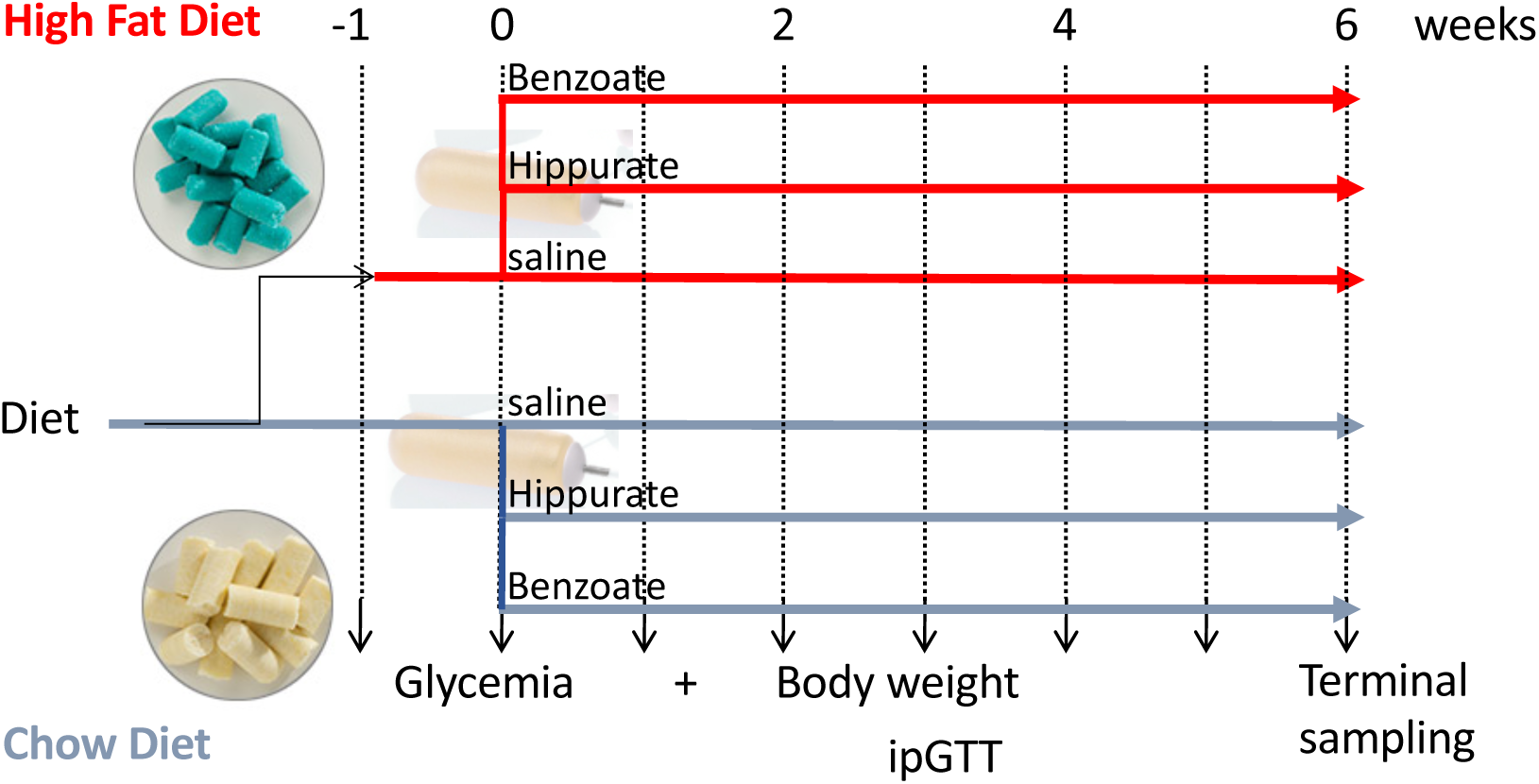
Experimental design for chronic six-week administration of benzoate and hippurate. Experiment design showing groups and durations of each step for the chronic treatments with benzoate and hippurate in mice.

**Supplementary Figure 4.**
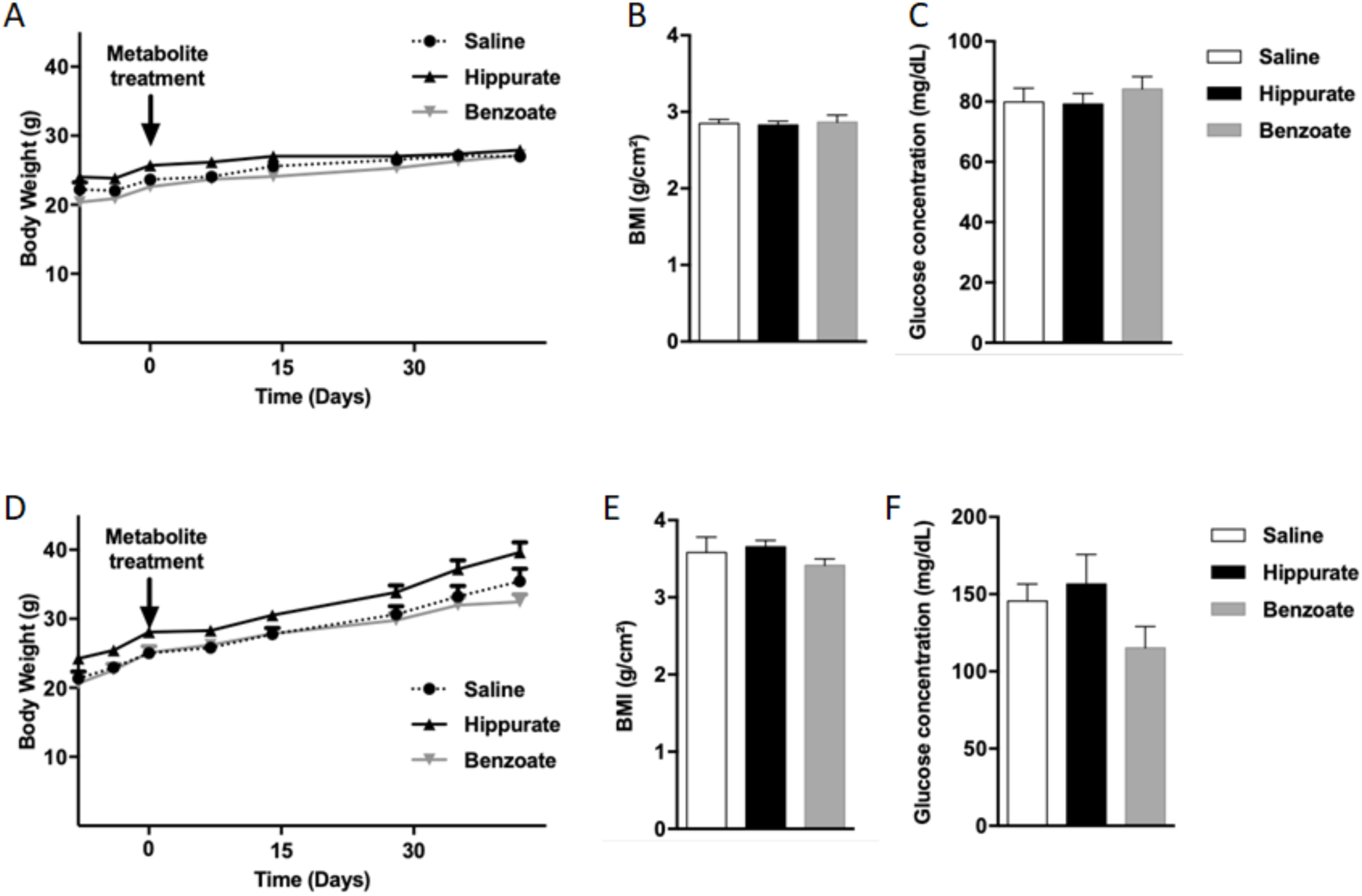
Effects of chronic administration of hippurate and benzoate on body growth and fasting glycemia. C57BL6/J mice fed control chow diet **(A-C)** or high fat diet **(D-F)**. The effects of chronic subcutaneous administration of the metabolites (5.55 mM) in mice were tested on body weight **(A,D)**, body mass index (BMI) **(B,E)**, fasting glycemia **(C,F)**. Control mice were treated with saline. BMI was calculated as body weight divided by the squared of anal-nasal length. Results are means ± SEM.

**Supplementary Figure 5.**
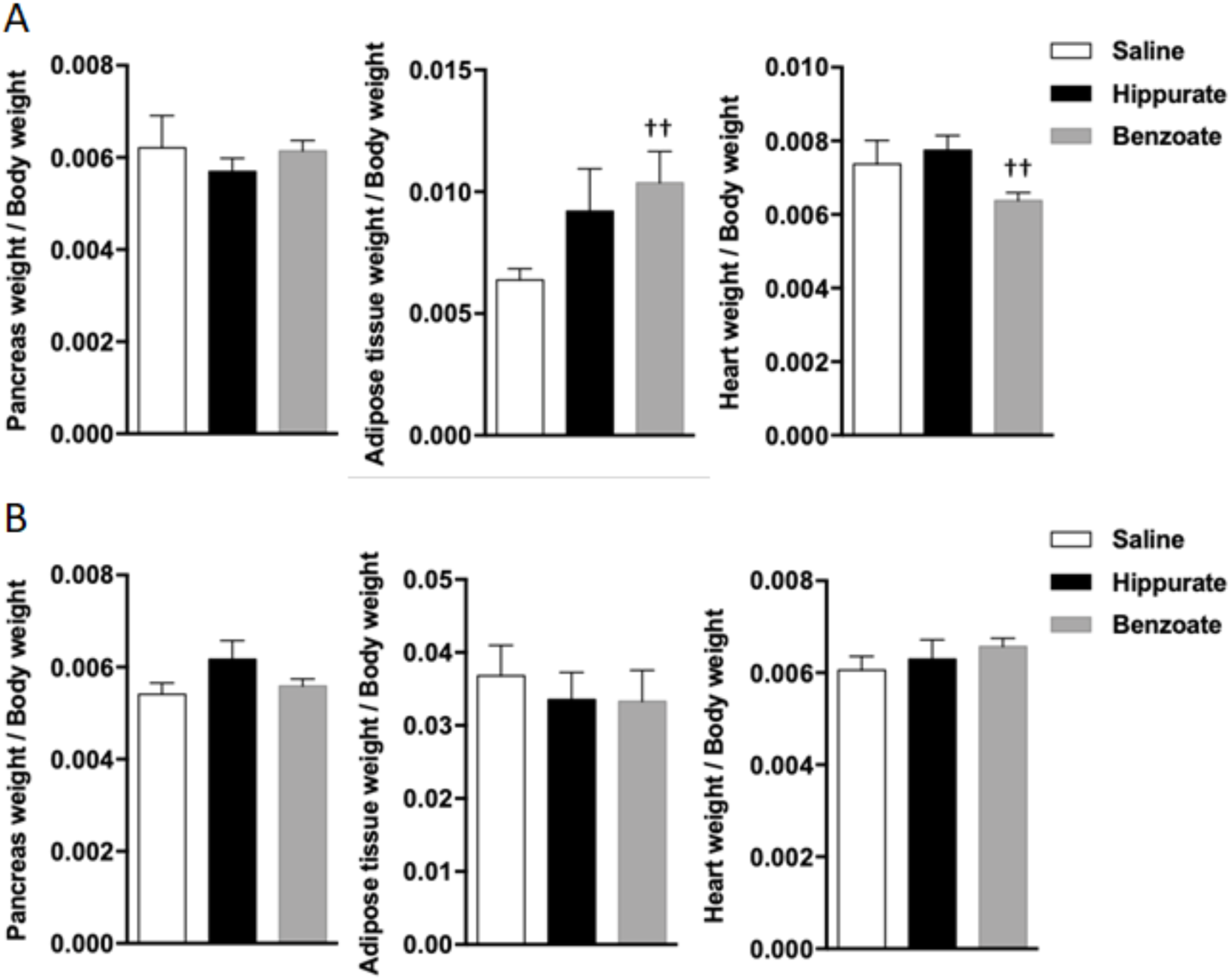
Organ weight in C57BL6/J mice treated chronically with hippurate or benzoate for 42 days. Control mice were treated with saline. Mice were fed chow diet (CHD) or high fat diet (HFD) for 56 days. Data are expressed as the ratio between organ weight and body weight. Data were analyzed using the unpaired Mann-Whitney test. Results are means ±SEM. **P<0.01, significantly different between mice treated with hippurate and controls. †P< 0.05, ††P<0.01, significantly different between mice treated with benzoate and saline treated controls.

**Supplementary Figure 6.**
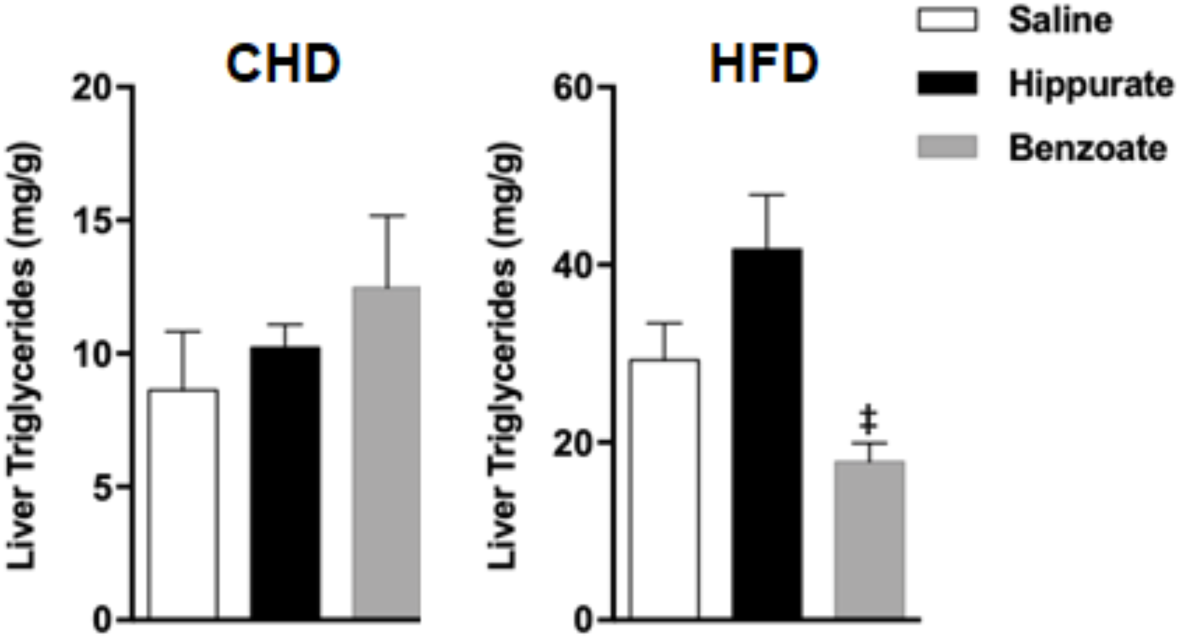
Effect of chronic administration of hippurate and benzoate on liver triglycerides content in C57BL6/J mice. The effect of chronic subcutaneous administration of the metabolites (5.55mM) for 42 days on liver triglycerides was tested in mice fed control chow diet (CHD) or high fat diet (HFD) for 56 days. Assay was carried out in 6 mice per group. Data were analyzed using the unpaired Mann-Whitney test. Results are means ± SEM ‡P<0.05, significantly different between mice treated with benzoate and hippurate.

**Supplementary Table S1**

Association between microbiota functional potential mapped to KEGG Orthologs (KOs) database and urine hippurate levels.

**Supplementary Table S2**

Association between microbiota functional potential mapped to the eggNOG database and urine hippurate levels.

**Supplementary Table S3**

Association between microbiota functional potential mapped to gut-specific metabolic modules (GMMs) describing phenylpropanoid metabolism and urine hippurate levels.

**Supplementary Table S4**

Association between abundance of gut-specific metabolic modules (GMMs) describing phenylpropanoid metabolism and gene richness.

**Supplementary Table S5**

Phenylpropanoid metabolism potential in metagenomic species, assessed by mapping to GMMs significantly associated to urine hippurate levels.

**Supplementary Table S6**

Association between the microbiota composition profiled as metagenomic OTUs (mOTUs) and urine hippurate levels.

**Supplementary Table S7**

pFDR from Spearman’s rank-based correlations between GMMs describing phenylpropanoid metabolism and gene richnessbioclinical variables using Storey’s FDR correction.

**Supplementary Table 8**

pFDR from Mann-Whitney U test for hippurate stratification in each bioclinical variable between using Storey’s FDR correction.

## Supplementary List: Module definitions

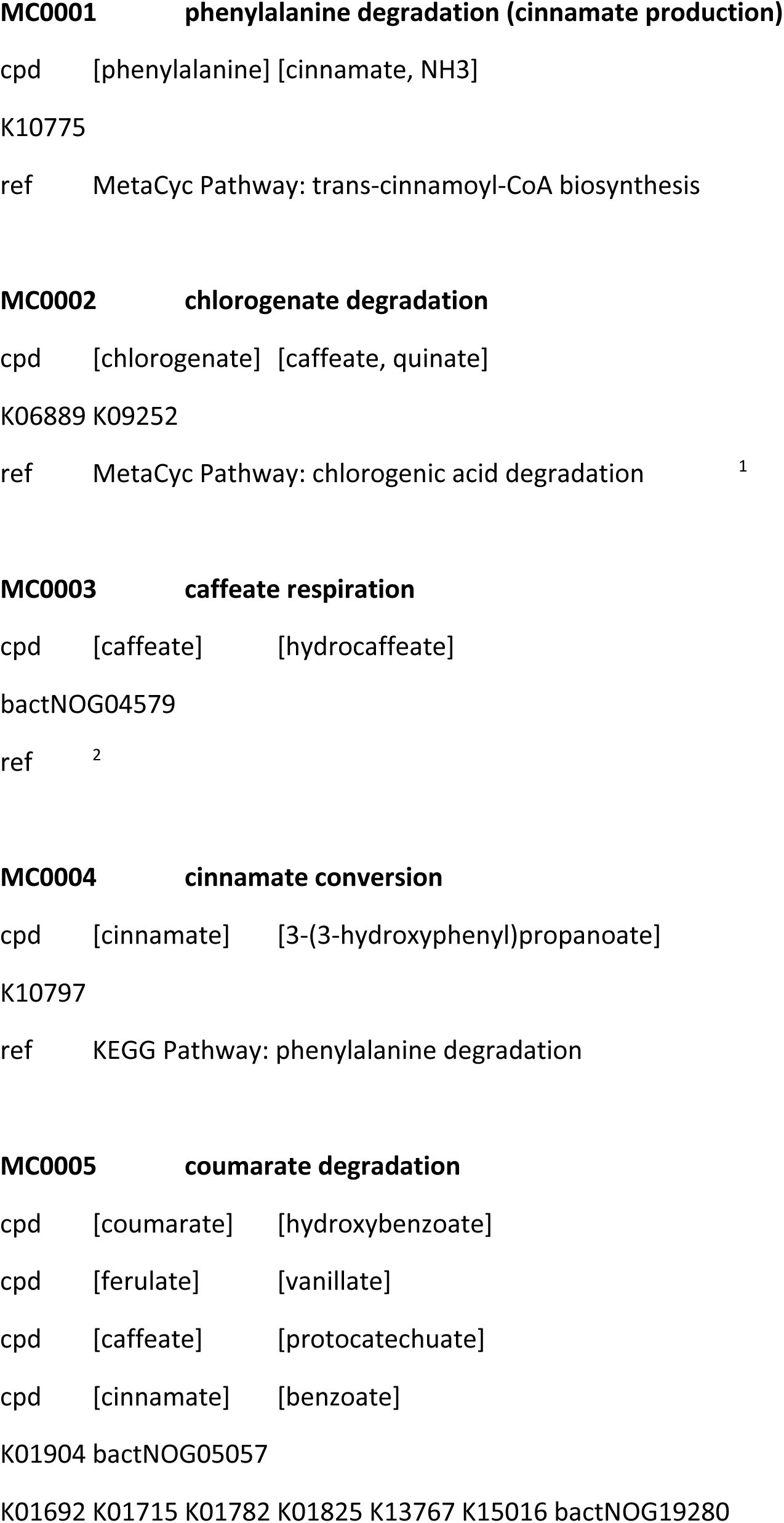
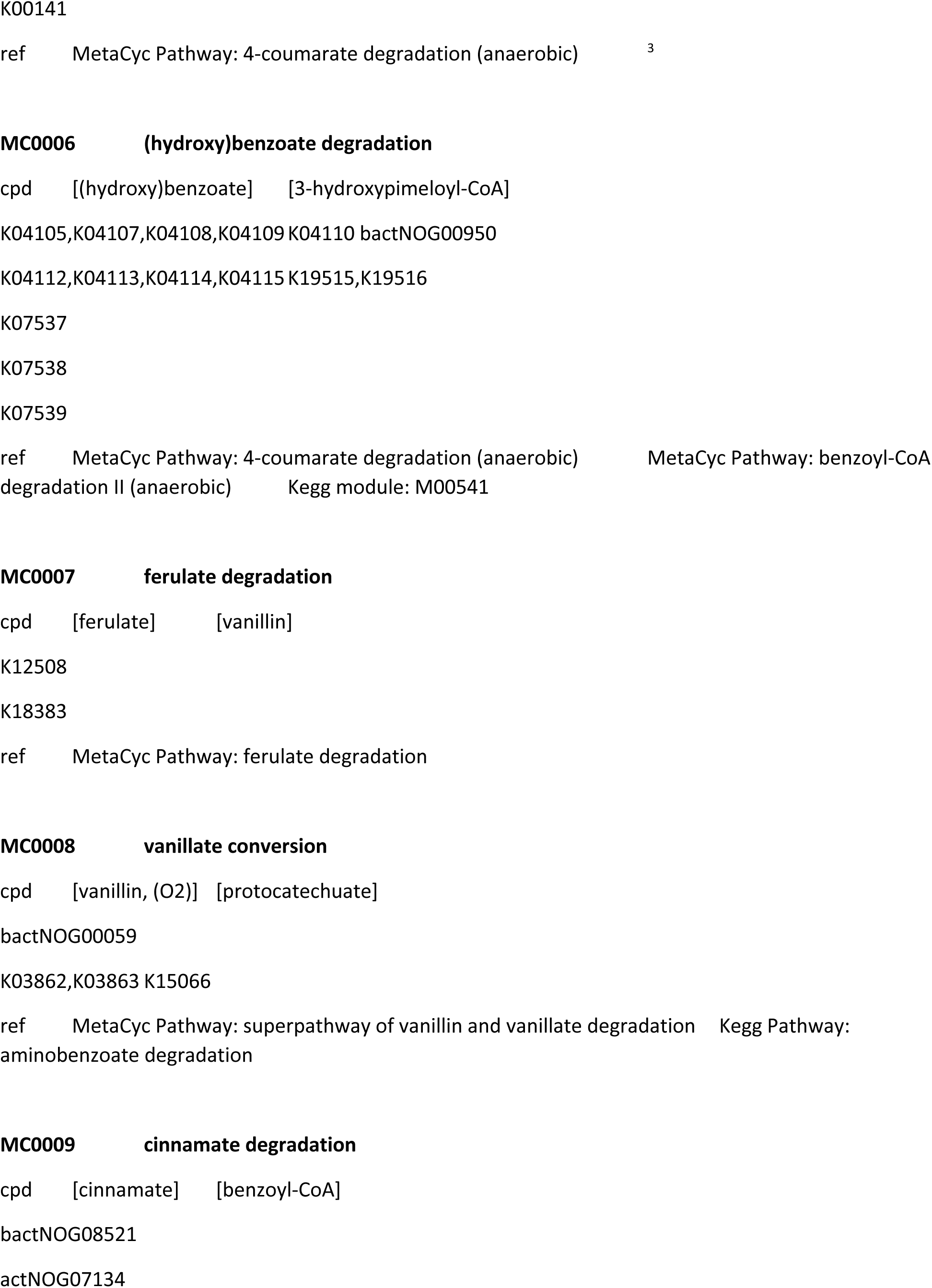
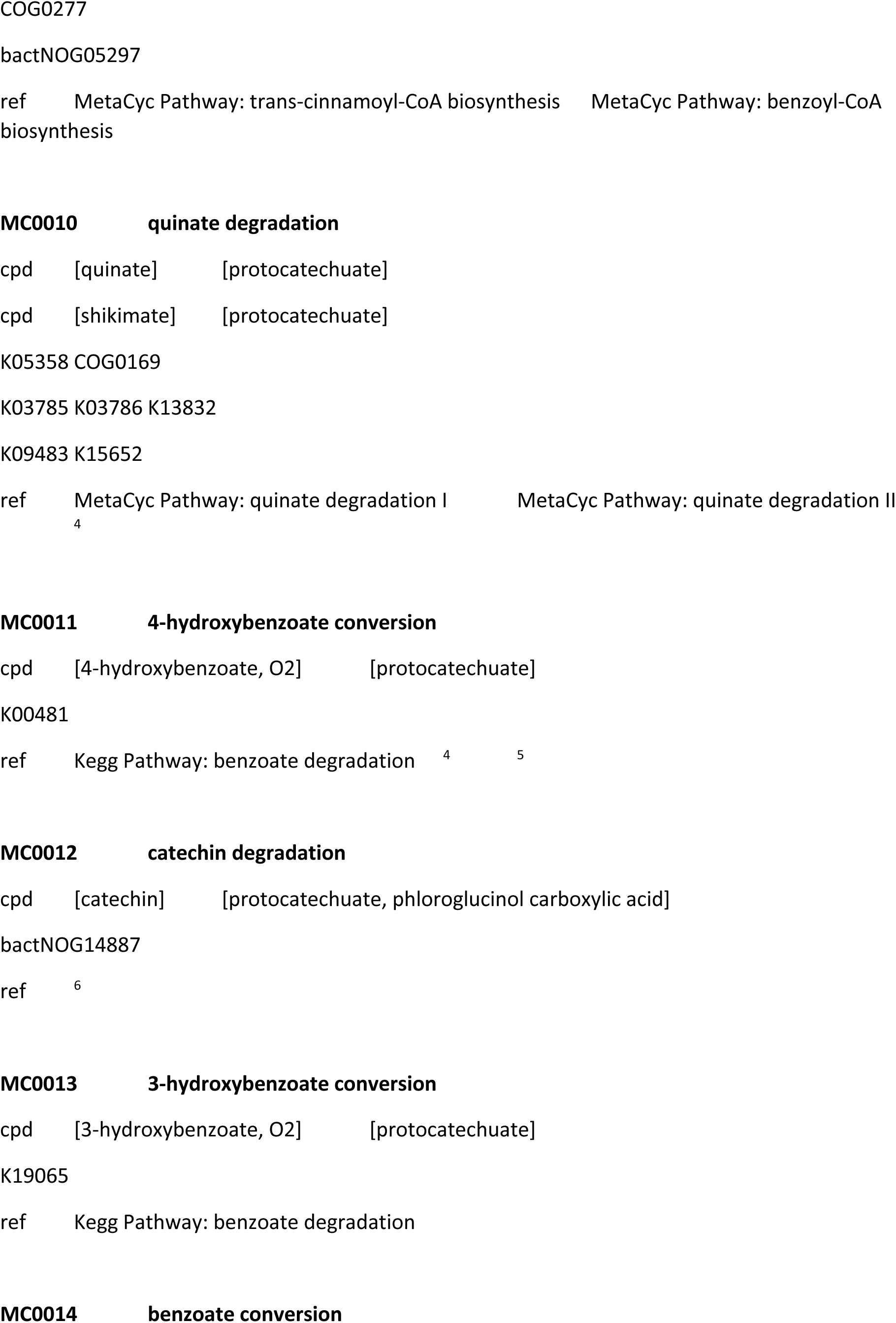
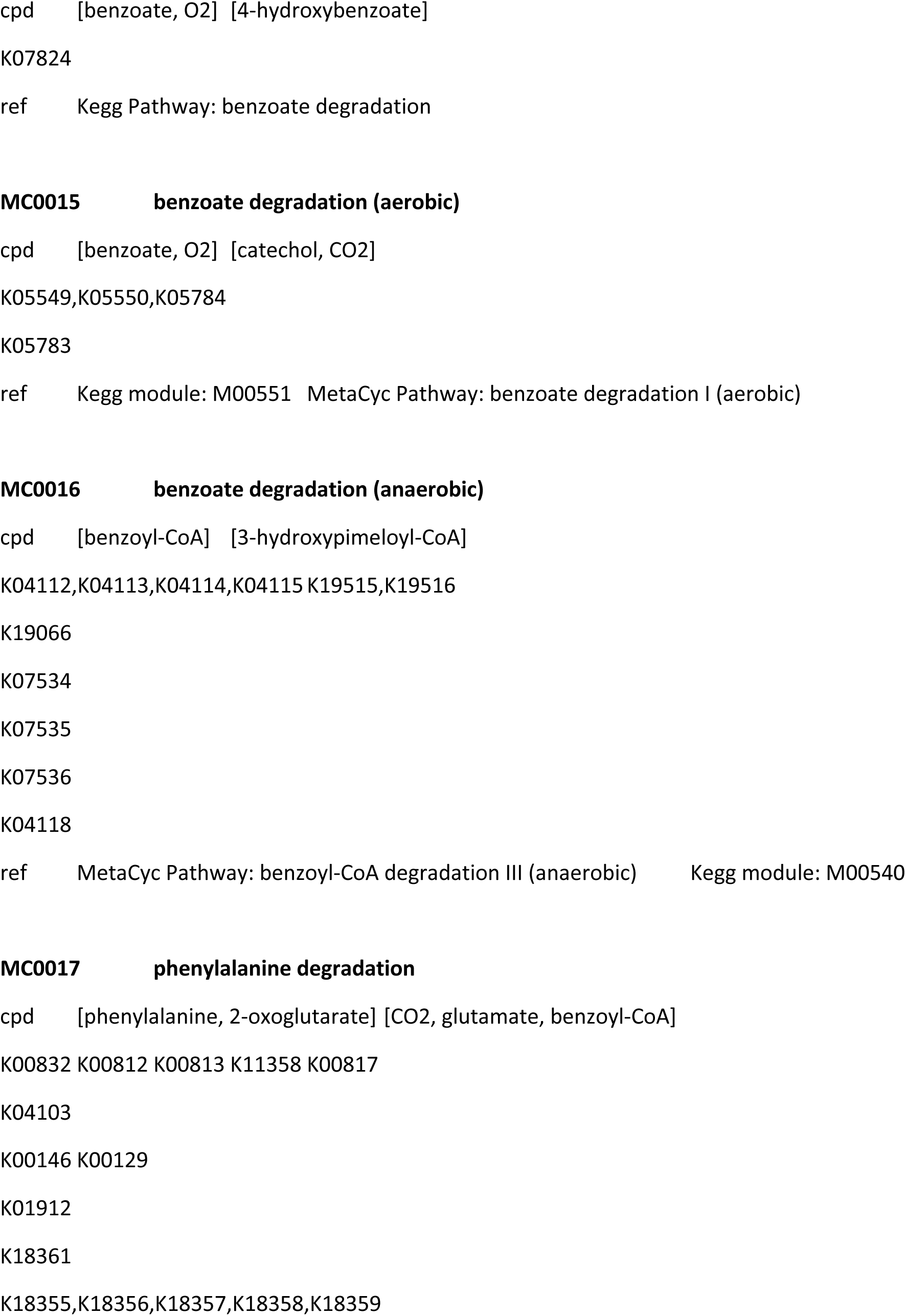
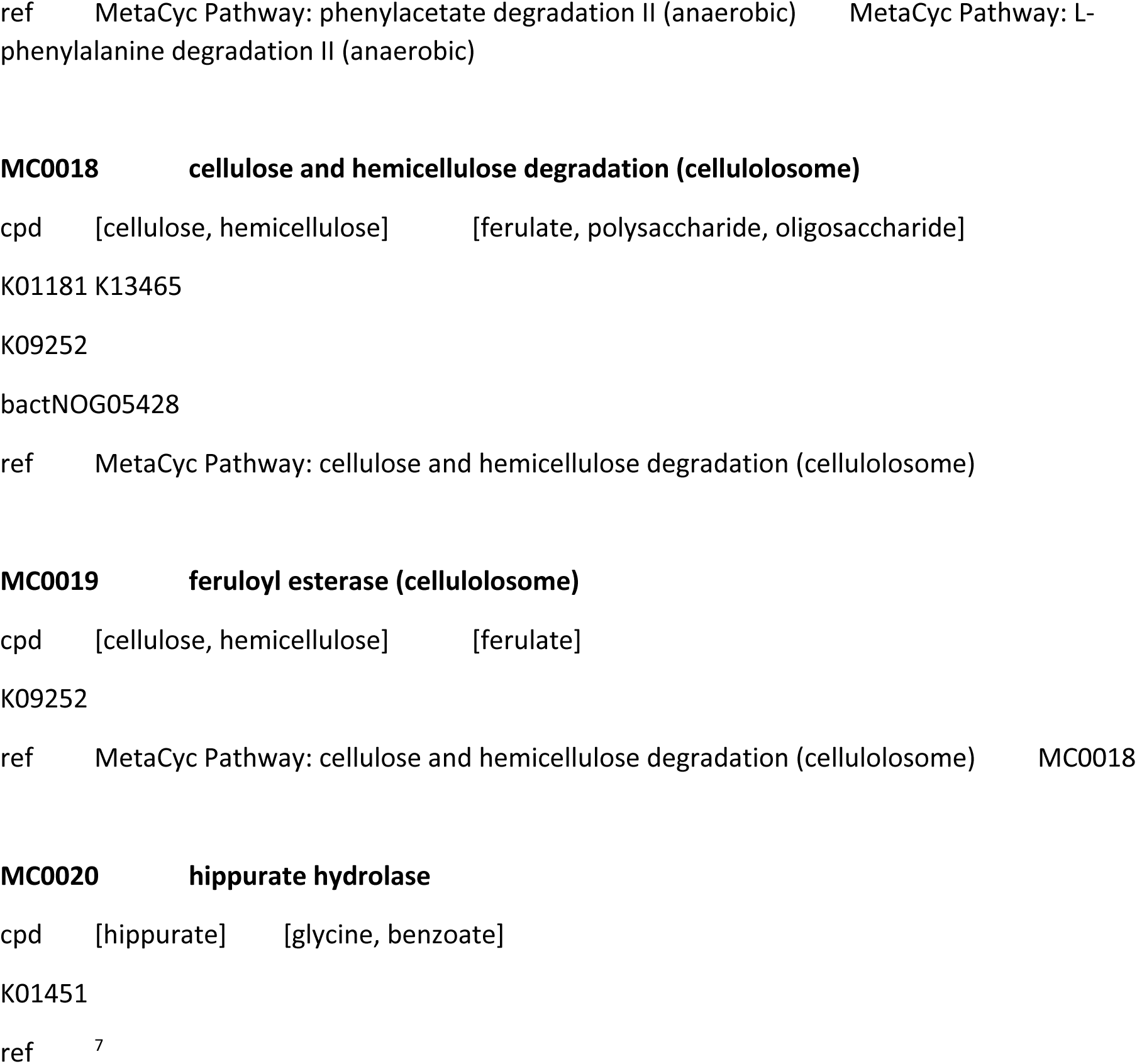

## REFERENCES

1. Lynch SV, Pedersen O. The Human Intestinal Microbiome in Health and Disease. N Engl J Med 2016;375:2369–79. doi:10.1056/NEJMra1600266

2. Li J, Jia H, Cai X, et al. An integrated catalog of reference genes in the human gut microbiome. Nat Biotechnol 2014;32:834–41. doi:10.1038/nbt.2942

3. Qin J, Li R, Raes J, et al. A human gut microbial gene catalogue established by metagenomic sequencing. Nature 2010;464:59–65. doi:10.1038/nature08821

4. Le Chatelier E, Nielsen T, Qin J, et al. Richness of human gut microbiome correlates with metabolic markers. Nature 2013;500:541–6. doi:10.1038/nature12506

5. Cotillard A, Kennedy SP, Kong LC, et al. Dietary intervention impact on gut microbial gene richness. Nature 2013;500:585–8. doi:10.1038/nature12480

6. Nicholson JK, Holmes E, Wilson ID. Gut microorganisms, mammalian metabolism and personalized health care. Nat Rev Microbiol 2005;3:431–8. doi:10.1038/nrmicro1152

7. Dumas M-E. The microbial-mammalian metabolic axis: beyond simple metabolism. Cell Metab 2011;13:489–90. doi:10.1016/j.cmet.2011.04.005

8. Nicholson JK, Holmes E, Kinross J, et al. Host-gut microbiota metabolic interactions. Science 2012;336:1262–7. doi:10.1126/science.1223813

9. Neves AL, Chilloux J, Sarafian MH, et al. The microbiome and its pharmacological targets: therapeutic avenues in cardiometabolic diseases. Curr Opin Pharmacol 2015;25:36–44. doi:10.1016/j.coph.2015.09.013

10. Dumas M-E, Barton RH, Toye A, et al. Metabolic profiling reveals a contribution of gut microbiota to fatty liver phenotype in insulin-resistant mice. Proc Natl Acad Sci USA 2006;103:12511–6. doi:10.1073/pnas.0601056103

11. Holmes E, Loo RL, Stamler J, et al. Human metabolic phenotype diversity and its association with diet and blood pressure. Nature 2008;453:396–400. doi:10.1038/nature06882

12. Elliott P, Posma JM, Chan Q, et al. Urinary metabolic signatures of human adiposity. Sci Transl Med 2015;7:285ra62. doi:10.1126/scitranslmed.aaa5680

13. Pallister T, Jackson MA, Martin TC, et al. Untangling the relationship between diet and visceral fat mass through blood metabolomics and gut microbiome profiling. Int J Obes (Lond) 2017;41:1106–13. doi:10.1038/ijo.2017.70

14. Hoyles L, Fernández-Real JM, Federici M, et al. Molecular phenomics and metagenomics of hepatic steatosis in non-diabetic obese women. Nat Med 2018;24:1070–80. doi:10.1038/s41591-018-0061-3

15. Lees HJ, Swann JR, Wilson ID, et al. Hippurate: The Natural History of a Mammalian-Microbial Cometabolite. J Proteome Res 2013;12:1527–46. doi:10.1021/pr300900b

16. Dumas M-E, Wilder SP, Bihoreau M-T, et al. Direct quantitative trait locus mapping of mammalian metabolic phenotypes in diabetic and normoglycemic rat models. Nat Genet 2007;39:666–72. doi:10.1038/ng2026

17. Pallister T, Jackson MA, Martin TC, et al. Hippurate as a metabolomic marker of gut microbiome diversity: Modulation by diet and relationship to metabolic syndrome. Scientific Reports 2017;7:13670. doi:10.1038/s41598-017-13722-4

18. Pedersen HK, Gudmundsdottir V, Nielsen HB, et al. Human gut microbes impact host serum metabolome and insulin sensitivity. Nature 2016;535:376–81. doi:10.1038/nature18646

19. Forslund K, Hildebrand F, Nielsen T, et al. Disentangling type 2 diabetes and metformin treatment signatures in the human gut microbiota. Nature 2015;528:262–6. doi:10.1038/nature15766

20. Jørgensen T, Borch-Johnsen K, Thomsen TF, et al. A randomized non-pharmacological intervention study for prevention of ischaemic heart disease: baseline results Inter99. Eur J Cardiovasc Prev Rehabil 2003;10:377–86. doi:10.1097/01.hjr.0000096541.30533.82

21. Levey AS, Stevens LA, Schmid CH, et al. A new equation to estimate glomerular filtration rate. Ann Intern Med 2009;150:604–12.

22. Toft U, Kristoffersen L, Ladelund S, et al. Relative validity of a food frequency questionnaire used in the Inter99 study. Eur J Clin Nutr 2008;62:1038–46. doi:10.1038/sj.ejcn.1602815

23. Dona AC, Jiménez B, Schäfer H, et al. Precision high-throughput proton NMR spectroscopy of human urine, serum, and plasma for large-scale metabolic phenotyping. Anal Chem 2014;86:9887–94. doi:10.1021/ac5025039

24. Blaise BJ, Shintu L, Elena B, et al. Statistical recoupling prior to significance testing in nuclear magnetic resonance based metabonomics. Anal Chem 2009;81:6242–51. doi:10.1021/ac9007754

25. Dona AC, Kyriakides M, Scott F, et al. A guide to the identification of metabolites in NMR-based metabonomics/metabolomics experiments. Comput Struct Biotechnol J 2016;14:135–53. doi:10.1016/j.csbj.2016.02.005

26. Vieira-Silva S, Falony G, Darzi Y, et al. Species-function relationships shape ecological properties of the human gut microbiome. Nat Microbiol 2016;1:16088. doi:10.1038/nmicrobiol.2016.88

27. Storey JD, Tibshirani R. Statistical significance for genomewide studies. Proc Natl Acad Sci USA 2003;100:9440–5. doi:10.1073/pnas.1530509100

28. Dixon P. VEGAN, a package of R functions for community ecology. Journal of Vegetation Science 2003;14:927–30. doi:10.1111/j.1654-1103.2003.tb02228.x

29. Cloarec O, Dumas ME, Trygg J, et al. Evaluation of the orthogonal projection on latent structure model limitations caused by chemical shift variability and improved visualization of biomarker changes in 1H NMR spectroscopic metabonomic studies. Anal Chem 2005;77:517–26. doi:10.1021/ac048803i

30. Blaise BJ, Giacomotto J, Elena B, et al. Metabotyping of Caenorhabditis elegans reveals latent phenotypes. Proc Natl Acad Sci USA 2007;104:19808–12. doi:10.1073/pnas.0707393104

31. Brial F, Le Lay A, Hedjazi L, et al. Systems Genetics of Hepatic Metabolome Reveals Octopamine as a Target for Non-Alcoholic Fatty Liver Disease Treatment. Scientific Reports 2019;9:3656. doi:10.1038/s41598-019-40153-0

32. Phipps AN, Stewart J, Wright B, et al. Effect of diet on the urinary excretion of hippuric acid and other dietary-derived aromatics in rat. A complex interaction between diet, gut microflora and substrate specificity. Xenobiotica 1998;28:527–37. doi:10.1080/004982598239443

33. Backhed F, Ding H, Wang T, et al. The gut microbiota as an environmental factor that regulates fat storage. Proc Natl Acad Sci USA 2004;101:15718–23. doi:10.1073/pnas.0407076101

34. Bridle KR, Crawford DHG, Ramm GA. Identification and characterization of the hepatic stellate cell transferrin receptor. Am J Pathol 2003;162:1661–7. doi:10.1016/S0002-9440(10)64300-3

35. Shi J, Zhao J, Zhang X, et al. Activated hepatic stellate cells impair NK cell anti-fibrosis capacity through a TGF-β-dependent emperipolesis in HBV cirrhotic patients. Scientific Reports 2017;7:44544. doi:10.1038/srep44544

36. Aron-Wisnewsky J, Prifti E, Belda E, et al. Major microbiota dysbiosis in severe obesity: fate after bariatric surgery. Gut 2019;68:70–82. doi:10.1136/gutjnl-2018-316103

37. Akira K, Masu S, Imachi M, et al. 1H NMR-based metabonomic analysis of urine from young spontaneously hypertensive rats. J Pharm Biomed Anal 2008;46:550–6. doi:10.1016/j.jpba.2007.11.017

38. Zhao L-C, Zhang X-D, Liao S-X, et al. A metabonomic comparison of urinary changes in Zucker and GK rats. J Biomed Biotechnol 2010;2010:431894–6. doi:10.1155/2010/431894

39. Pontoizeau C, Fearnside JF, Nayratil V, et al. Broad-Ranging Natural Metabotype Variation Drives Physiological Plasticity in Healthy Control Inbred Rat Strains. J Proteome Res 2011;10:1675–89. doi:10.1021/pr101000z

40. Bitner BF, Ray JD, Kener KB, et al. Common gut microbial metabolites of dietary flavonoids exert potent protective activities in β-cells and skeletal muscle cells. J Nutr Biochem 2018;62:95–107. doi:10.1016/j.jnutbio.2018.09.004

41. Dumas M-E, Rothwell AR, Hoyles L, et al. Microbial-Host Co-metabolites Are Prodromal Markers Predicting Phenotypic Heterogeneity in Behavior, Obesity, and Impaired Glucose Tolerance. Cell Rep 2017;20:136–48. doi:10.1016/j.celrep.2017.06.039

42. Thaiss CA, Itav S, Rothschild D, et al. Persistent microbiome alterations modulate the rate of post-dieting weight regain. Nature 2016;540:540–51. doi:10.1038/nature20796

43. Patterson AD, Turnbaugh PJ. Microbial determinants of biochemical individuality and their impact on toxicology and pharmacology. Cell Metab 2014;20:761–8. doi:10.1016/j.cmet.2014.07.002

44. Shoaie S, Ghaffari P, Kovatcheva-Datchary P, et al. Quantifying Diet-Induced Metabolic Changes of the Human Gut Microbiome. Cell Metab 2015;22:320–31. doi:10.1016/j.cmet.2015.07.001

45. Zeevi D, Korem T, Zmora N, et al. Personalized Nutrition by Prediction of Glycemic Responses. Cell 2015;163:1079–94. doi:10.1016/j.cell.2015.11.001

## References

1. Couteau, D., McCartney, A. L., Gibson, G. R., Williamson, G. & Faulds, C. B. Isolation and characterization of human colonic bacteria able to hydrolyse chlorogenic acid. J. Appl. Microbiol. 90, 873–881 (2001).

2. Hess, V., González, J. M., Parthasarathy, A., Buckel, W. & Müller, V. Caffeate respiration in the acetogenic bacterium Acetobacterium woodii: a coenzyme A loop saves energy for caffeate activation. Appl. Environ. Microbiol. 79, 1942–1947 (2013).

3. Hirakawa, H., Schaefer, A. L., Greenberg, E. P. & Harwood, C. S. Anaerobic p-coumarate degradation by Rhodopseudomonas palustris and identification of CouR, a MarR repressor protein that binds p-coumaroyl coenzyme A. J. Bacteriol. 194, 1960–1967 (2012).

4. Brzostowicz, P. C., Reams, A. B., Clark, T. J. & Neidle, E. L. Transcriptional cross-regulation of the catechol and protocatechuate branches of the beta-ketoadipate pathway contributes to carbon source-dependent expression of the Acinetobacter sp. strain ADP1 pobA gene. Appl. Environ. Microbiol. 69, 1598–1606 (2003).

5. Gonthier, M.-P. et al. Metabolism of dietary procyanidins in rats. Free Radic. Biol. Med. 35, 837–844 (2003).

6. Arunachalam, M., Mohan, N. & Mahadevan, A. Cloning of Acinetobacter calcoaceticus chromosomal region involved in catechin degradation. Microbiol. Res. 158, 37–46 (2003).

7. Caner, V., Cokal, Y., Cetin, C., Sen, A. & Karagenc, N. The detection of hipO gene by real-time PCR in thermophilic Campylobacter spp. with very weak and negative reaction of hippurate hydrolysis. Antonie Van Leeuwenhoek 94, 527–532 (2008).

